# The roles and evolution of the four *LEAFY* homologues in floral patterning and leaf development in woodland strawberry

**DOI:** 10.1101/2022.10.06.511044

**Authors:** Yunming Zhang, Lijun Kan, Shaoqiang Hu, Laichao Cheng, Zhongchi Liu, Chunying Kang

## Abstract

The plant-specific transcription factor *LEAFY* (*LFY*), generally maintained as a single copy gene in most angiosperm species, plays critical roles in flower and leaf development. However, wild strawberry *Fragaria vesca* possesses four *LFY* homologues in the genome, their respective functions and evolution remain unknown. Through chemical mutagenesis screen, we identified two allelic mutations in one of the four *LFY* homologues, *FveLFYa*, in *F. vesca*, causing homeotic conversion of floral organs and reiterative outgrowth of ectopic florets. Both CRISPR-knockout and transgenic rescue confirmed the identity of *FveLFYa*. Ectopic expression in *Arabidopsis lfy-5* mutant revealed that only *FveLFYa* and *FveLFYb* can rescue the flower defects and induce solitary flowers in leaf axils. Disruption of *FveLFYc*, the second abundantly expressed *LFY* homologue, caused no obvious morphology phenotypes in *F. vesca. FveLFYb* and *FveLFYd* are barely expressed. Expression of *FveAP1*, homologue of the well-known LFY target *AtAP1*, is not changed in the *fvelfya* flowers, possibly caused by an absence of any FveLFYa binding site in its promoter. *Loss of Axillary Meristems* encodes a GRAS transcription factor essential for stamen initiation. The ectopic florets are eliminated in *fvelfya lam*, suggesting that *LAM* is required for floret production. Moreover, approximately 30% of mature leaves have smaller or fewer leaflets in *fvelfya*. Among these homologues, only *FveLFYa* is syntenic to the homologues in other species. Overall, the detailed analyses of the four *LFY* homologues in woodland strawberry demonstrate that only *FveLFYa* plays crucial roles in floral patterning with rewired gene network.

## Introduction

LEAFY (LFY) is a highly conserved transcription factor that exists in all land plants and algae (Sayou et al., 2014; Gao et al., 2019). *LFY* plays versatile roles from cell division to plant development, especially in flowering and floral patterning (Moyroud et al., 2009; Moyroud et al., 2010). The rice *LFY* homolog *RFL* is positive regulator of flowering time (Rao et al., 2008). Overexpression of *LFY* converted the lateral shoots to solitary flowers in diverse plants, indicating it is sufficient for flower initiation (Weigel and Nilsson, 1995). Of note, the *lfy* mutants show variable flower morphologies in different angiosperm species (Coen et al., 1990; Weigel et al., 1992; Wang et al., 2008; Monniaux et al., 2017). *LFY* disruption causes the complete homeotic conversion from a flower to a shoot in *Antirrhinum majus* and *Cardamine hirsute* (Coen et al., 1990; Monniaux et al., 2017). The *lfy* strong allele shows only a partial homeotic conversion of flowers into leafy shoots in *Arabidopsis* (Schultz and Haughn, 1991; Weigel et al., 1992). The *Medicago truncatula LFY* mutant *sgl1* produce cauliflower-like flowers (Wang et al., 2008). These findings emphasize the value of investigating *LFY* in different species to fully uncover the mechanism of this important regulator of flowers.

*LFY* is involved in regulating leaf development in some species. In the inverted-repeat-lacking clade (IRLC) legumes, mutation of the *LFY* orthologues *UNIFOLIATA* (*UNI*) in *P. sativum* and *SGL1* in *M. truncatula* results in a transition from compound leaves to simple leaves (Hofer et al., 1997; Wang et al., 2008). Other *lfy* mutants show a reduced number of leaflets on the compound leaves in tomato, *C. hirsuta* and some non-IRLC legumes (Molinero-Rosales et al., 1999; Champagne et al., 2007; Monniaux et al., 2017; Jiao et al., 2019). In *Arabidopsis*, although no leaf phenotype was observed in *lfy* mutants, *LFY* is required for the ruffled leaf margin in the *UNUSUAL FLORAL ORGANS* (*UFO*) overexpressed plants (Lee et al., 1997). Therefore, *LFY* probably plays a role in leaf development in more angiosperm lineages.

*LFY* orchestrates the gene regulatory network during flower development. In *A. thaliana*, LFY directly activates the A class gene *APETALA1* (*AP1*), creating a positive feedback loop to trigger floral meristem formation (Parcy et al., 1998; Wagner et al., 1999). The transcriptional links between LFY and *AP1* promoter is conserved in *C. hirsuta* (Monniaux et al., 2017). Subsequently, *LFY* activates the B and C class genes *APETALA3* (*AP3*), *PISTILLATA* (*PI*) and *AGAMOUS* (*AG*), in conjunction with the coregulators *UFO* and *WUSCHEL*, respectively (Lee et al., 1997; Parcy et al., 1998; Lenhard et al., 2001; Lohmann et al., 2001). Although the LFY binding sites in the large introns of *AG* homologues vary in number, position, sequence and affinity, the LFY/*AG* transcriptional link is well conserved in distant angiosperms species (Moyroud et al., 2011).

Unlike most transcription factors that form multigene families, *LFY* exists at a very low copy number, mostly as a single copy gene (Maizel et al., 2005; Sayou et al., 2014; Gao et al., 2019). In most angiosperms, *LFY* genes convergently reverted to single copy status after recurrent paleo-polyploidy duplication events (Van de Peer et al., 2017). The loss of *LFY* genes appears to be gene specific, rather than with large genomic regions (Gao et al., 2019). In species with recent duplications, additional copies of *LFY* exist, but seem not to be divergent in function (Wada et al., 2002; Bomblies et al., 2003). There are four *LFY* homologues in the woodland strawberry (*F. vesca*), a model species for the cultivated strawberry in Rosaceae. However, the *F. vesca* genome lacks the large-scale genome duplications seen in other rosids (Shulaev et al., 2011; Qiao et al., 2021). The date of the *LFY* gene duplications and fate of the duplicated genes remain unknown.

A typical *F. vesca* flower, from the outmost whorl to the innermost whorl, contains five bracts and five sepals, five petals, 20 stamens and more than 100 unfused carpels that are spirally arranged on a receptacle (Hollender et al., 2012). Here, we identified two EMS mutants Y535 and Y904 in *F. vesca*, showing homeotic conversion of petals, stamens, and partial carpels into sepals or bract-like organs, and iterative outgrowth of ectopic florets in each flower. These mutants are caused by disruption of the abundantly expressed *FveLFYa*. Further analyses revealed that *FveLFYa* is the predominant *LFY* gene and has lost the control of *FveAP1* expression. These results provide more insights into the functional divergence and evolution of the *LFY* homologues.

## RESULTS

### Isolation of mutants with secondary flowers iteratively formed at sepal axils in *F. vesca*

To identify floral regulators in strawberries, we screened an ethyl-methanesulfonate (EMS) mutagenized *F. vesca* (Yellow Wonder 5AF7, YW5AF7) population for floral patterning defects and isolated two mutants, Y535 and Y904. In these mutants, ectopic florets arise reiteratively at the sepal axils up to the quaternary level (Figure 1A, B). By contrast to five bracts and five sepals in a typical wild-type flower, approximately 40 sepals or sepal-like organs are generated in each Y535 or Y904 flower. The sepals or sepal-like organs in Y904 are wider than those of Y535 (Figure 1A, B). While virtually no petals are found in the Y535 flower, some petaloid organs can form in the primary flower of Y904 (Figure 1B). Stamens are completely lost in the flowers of both mutants. To determine whether Y535 and Y904 are mutants of the same causative gene, Y904/+ was crossed with Y535. Of the 17 progeny, eight plants show floral defects similar to Y904, and nine plants look like wild type, fitting to a 1:1 ratio (chi-square test, P = 0.81) (Supplemental Figure S1). This result suggests that Y535 and Y904 are allelic to each other.

**Figure 1.**
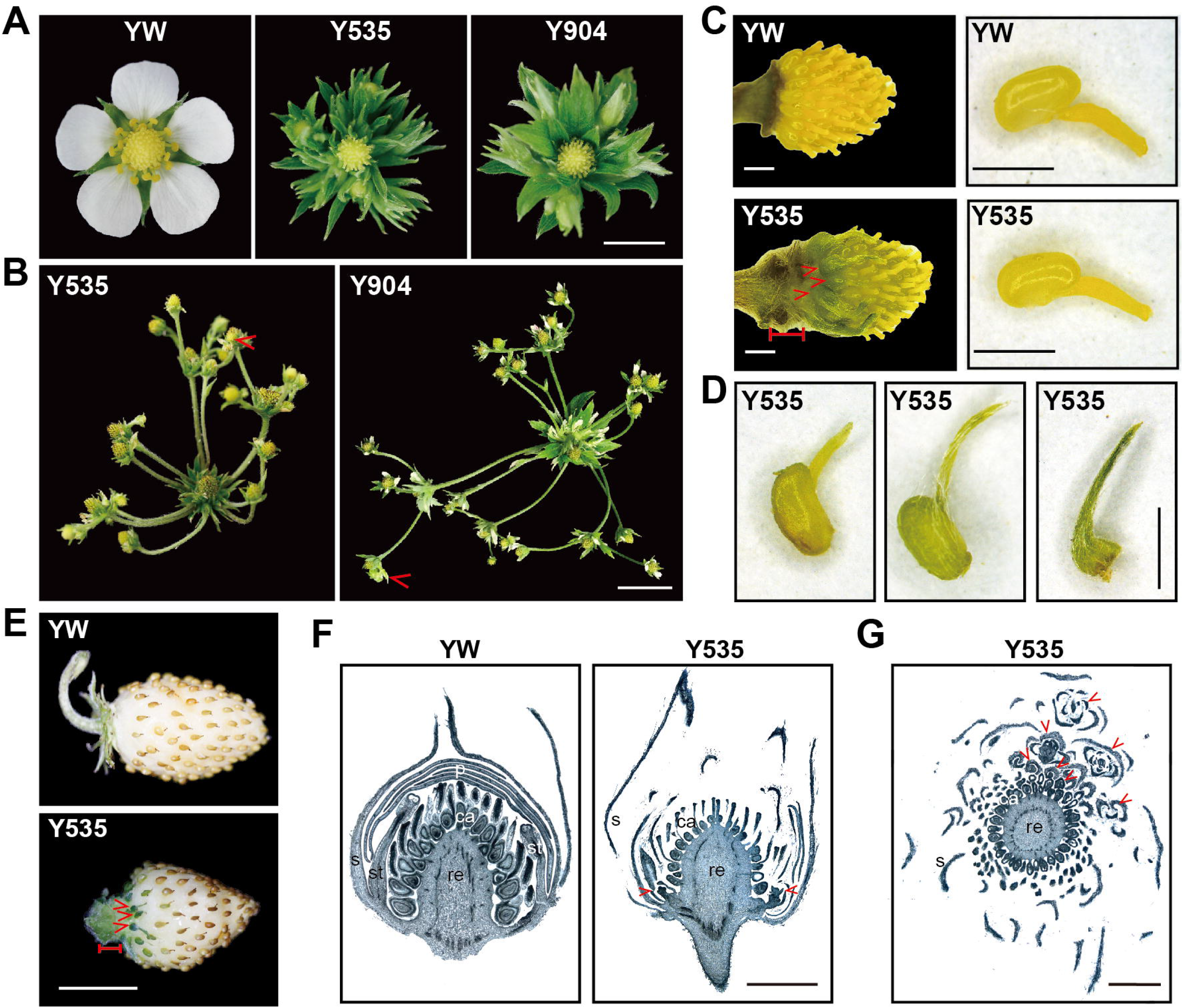
Phenotypic characterization of the *F. vesca* mutants Y535 and Y904. A, Flowers of YW, Y535 and Y904. B, Fully developed flowers of Y535 and Y904. C, The carpels on the receptacle and a single normal carpel of YW and Y535. D, Abnormal carpels of Y535. E, Mature fruit of YW and Y535. F. Longitudinal sections of YW and Y535 flower buds at stage 11. G. Transverse section of an Y535 flower bud at stage 11. s, sepal; p, petal; st, stamen; ca, carpel; re, receptacle. Red bars indicate the elongated internodes (C, E). Red arrowheads indicate the quaternary flowers (B), abnormal carpels (C, E), and secondary flowers (F, G). Scale bars: 0.5 cm (A), 1 cm (B, E), 1 mm (C-D, F, G).

We further characterized Y535’s floral defects. While all the wild type carpels are yellow, smooth, and have a fused indentation at the style apex (Figure 1C), some Y535 carpels at the base of the receptacle tend to be bract-like (Figure 1C, D). The bract-like organs are green, with epidermal hairs on the surface and sharp apexes, and sterile (Figure 1C-E). In addition, the internodes between sepals and the bottom carpels are elongated in Y535 flowers relative to WT flowers (Figure 1C, E). To observe the formation of secondary flowers, we used paraffin sections to visualize interior and higher order floral buds. Secondary flower primordia were observed at the sepal axils in Y535 flowers at stage 11 (Figure 1F). Interestingly, at every Y535 sepal axil, a secondary flower almost always forms (Figure 1G). Thus, the causative gene of Y535 and Y904 is required for floral organ specification, especially of petals and stamens, and to suppress the axillary meristem activity at their axils.

### The *LFY* homologue *FveLFYa* is the causative gene of Y535 and Y904

To determine the molecular basis of the mutants’ phenotype, we constructed F_2_ populations by crossing YW with Y535 and Y904, respectively. Both Y535 and Y904 are fully recessive as the F_1_ plants are indistinguishable from WT. Approximately one-quarter of Y535 (47 mutants out of 217 total plants) and Y904 (32 mutants out of 142 total plants) F_2_ plants exhibited the mutant phenotype, indicating that they are both caused by a recessive mutation in a single locus. Mapping-by-sequencing revealed that Y535 and Y904 each carried a different single nucleotide change in FvH4_5g09660, which encodes an LFY homologue and thus is named as *FveLFYa*. Y535 carries a C-to-T transition, generating a premature stop codon and resulting in a truncated protein composed of 37 amino acids without any conserved domain (Figure 2A). Y904 contains a G-to-A transition, causing an amino acid substitution of glutamate to lysine at position 114 in the conserved N-terminal domain (Figure 2A). The mutated glutamate is highly conserved among the LFY homologues (Supplemental Figure S2). Hereafter, we refer to Y535 and Y904 as *fvelfya-1* and *fvelfya-2*, respectively.

**Figure 2.**
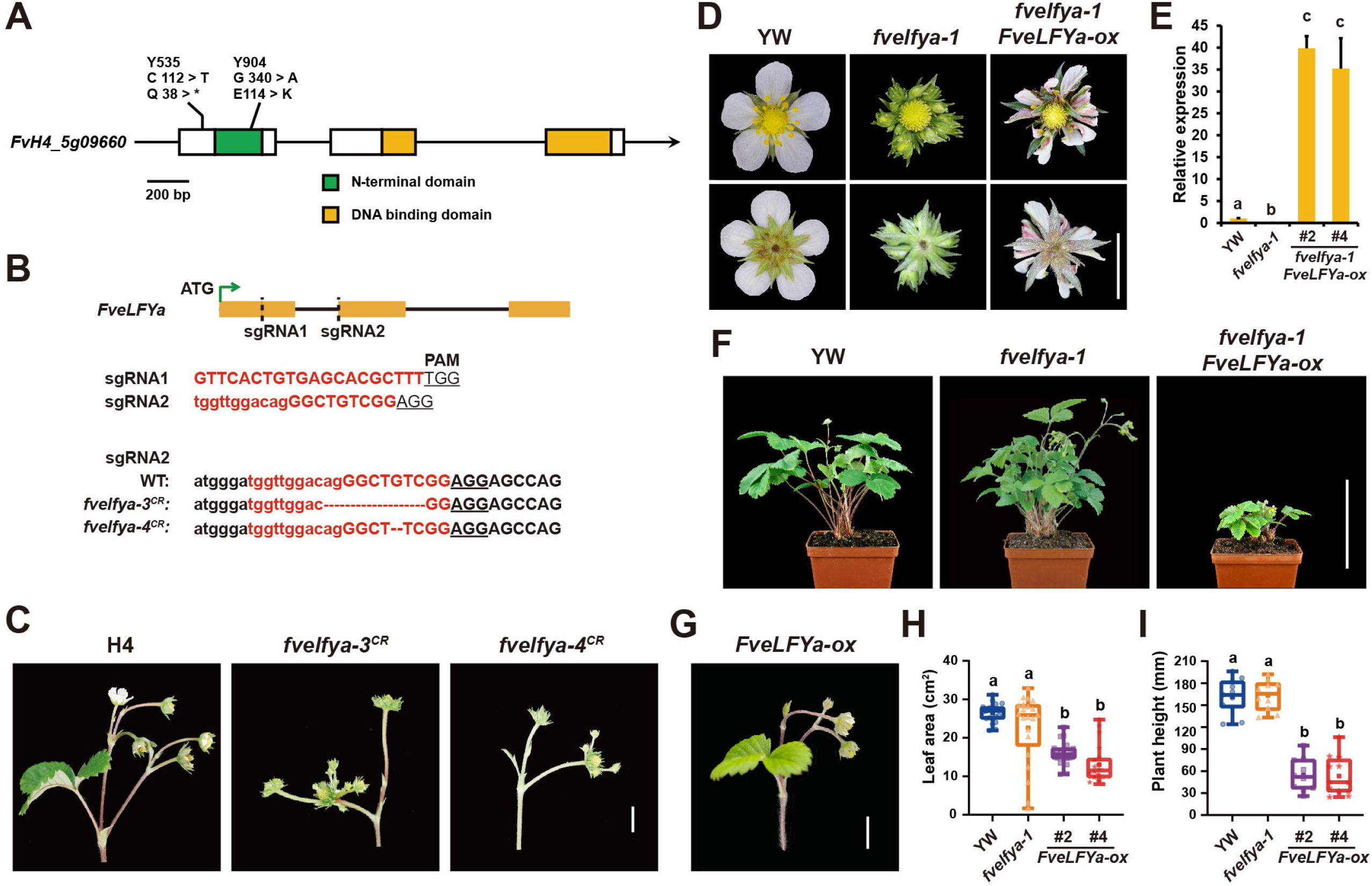
*FveLFYa* is the causative gene of Y535 and Y904. A, Schematic representation of the *FveLFYa* gene structure and the introduced mutations in Y535 (*fvelfya-1*) and Y904 (*fvelfya-2*). Rectangles represent exons; regions encoding the N-terminal domain (green) and the DNA-binding domain (yellow) are indicated. B, CRISPR-Cas9 targeting of *FveLFYa*. Deletions are indicated by dashes. Lowercase letters indicate intron sequence. Uppercase letters indicate exon sequence. C, Inflorescences of H4, *fvelfya-3*^*CR*^ and *fvelfya-4* ^*CR*^. D, Top and bottom views of the YW, *fvelfya-1* and *fvelfya-1 FveLFYa-ox* flowers. E, Relative expression levels of *FveLFYa* in flower buds at stages 3-7 of YW, *fvelfya-1* and *fvelfya-1 FveLFYa-ox* (#2 and #4). Data are the mean ±SD of three biological replicates. F, Side view of YW, *fvelfya-1* and *fvelfya-1 FveLFYa-ox* plants. G, An inflorescence of *FveLFYa-ox*. H and I, Box plots of leaf area (H) and plant height (I) of YW, *fvelfya-1* and *FveLFYa-ox* plants (n ≥ 10). Different letters indicate significant differences among plant lines (P < 0.01; one-way ANOVA and Tukey HSD). Scale bars: 1 cm (C, D, G); 10 cm (F).

To further confirm the identity of *FveLFYa*, we generated two additional *fvelfya* mutant alleles in the variety Hawaii-4 (H4) using CRISPR/Cas9. *fvelfya-3*^*CR*^ has a 9-nucleotide deletion (2 nucleotides located in the intron and 7 nucleotides located in the exon) at the sgRNA2 target site in *FveLFYa*, which may disrupt splicing and result in a protein without the DNA binding domain (Figure 2B). *fvelfya-4* ^*CR*^ has a G deletion in exon 2, which causes a frameshift leading to a stop codon after 186 amino acids. Hence, both *fvelfya-3* ^*CR*^ and *fvelfya-4* ^*CR*^ alleles are likely to be null. Their flowers also lost the petals and stamens, and generated ectopic florets, similar to those of *fvelfya-1* and *fvelfya-2* (Figure 2C). Furthermore, we overexpressed *FveLFYa* under the control of the cauliflower mosaic virus (CaMV) 35S promoter (*FveLFYa-ox*). The *fvelfya-1 FveLFYa-ox* flowers produce mosaic petals with sepal-like green tissues and a few intact stamens, and no secondary florets formed (Figure 2D). Expression levels of *FveLFYa* are indeed remarkably increased in the *fvelfya-1 FveLFYa-ox* leaves examined by qRT-PCR (Figure 2E). These results indicate that the floral defects in *fvelfya-1* are mostly rescued by the *FveLFYa* overexpression. In addition, ectopic expression of *FveLFYa* inhibits the growth of the entire plant. Compared to YW and *fvelfya-1*, the *FveLFYa-ox* plants showed reduced leaf size and decreased plant height (Figure 2F, H, I). In addition, the inflorescence structures of *FveLFYa-ox* are basically the same as those of H4 (wild-type), *fvelfya-3*^*CR*^, and *fvelfya-4*^*CR*^ (Figure 2C, G), suggesting that the formation of flower meristem is not altered by *FveLFYa*. Together, *FveLFYa* is an essential gene during floral patterning in *F. vesca* with roles in specifying proper floral organs and suppressing axillary meristems.

### FveLFYa and FveLFYb are functionally conserved AtLFY homologues

In addition to *FveLFYa*, there are three other *LFY* homologues in the *F. vesca* genome, designated as *FveLFYb* (FvH4_3g04170), *FveLFYc* (FvH4_3g03530) and *FveLFYd* (FvH4_3g03570), respectively. Of these, *FveLFYa*–*c* share the well-conserved *LFY* gene structure with three exons (Gao et al., 2019), while *FveLFYd* contains only the second and third exons of a typical *LFY* (Supplemental Figure S3). Consequently, the FveLFYa–c proteins contain the conserved N-terminal domain and C-terminal DNA binding domain, in contrast to the short sequence of FveLFYd (Figure 3A). Based on the DNA binding specificity, LFY can be classified into type-I, II, III and promiscuous LFY (Sayou et al., 2014). All the identified LFY proteins of angiosperms are type I LFY containing the three conserved amino acids (H312, R345, H387), and are supposed to be functionally conserved with AtLFY (Maizel et al., 2005; Sayou et al., 2014; Gao et al., 2019). We found that FveLFYa and FveLFYb have these conserved amino acids, so they should be the type I LFY. In FveLFYc, the residue R345 is replaced by methionine (M), suggesting that FveLFYc is atypical and may possess altered DNA binding property or play divergent functions.

**Figure 3.**
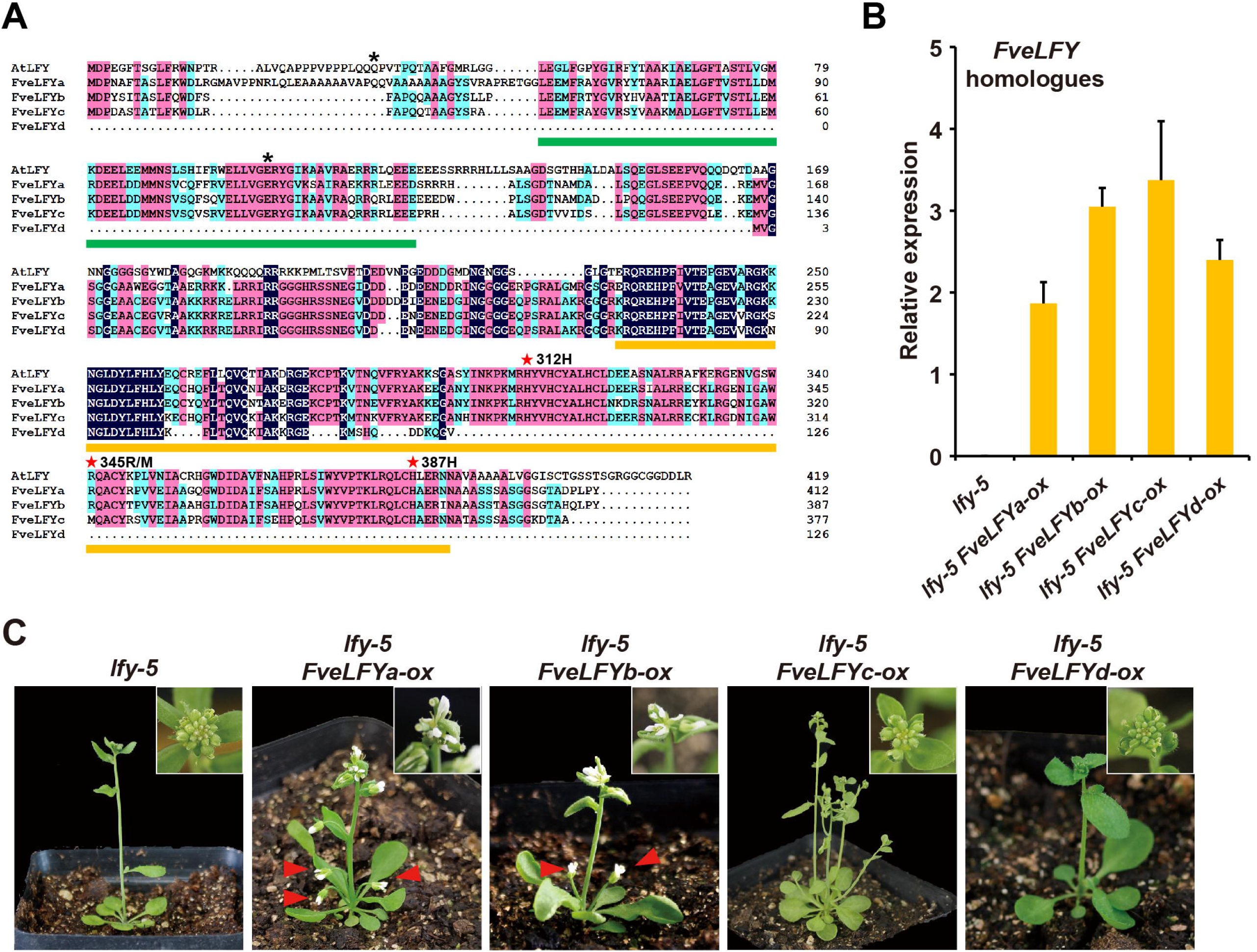
FveLFYa-d sequence alignment and functions in *Arabidopsis*. A, Amino acid sequence alignment of AtLFY and FveLFYs. The N terminal domain and DNA binding domain are labelled with green and yellow bottom lines, respectively. Asterisks indicate the mutation sites in *fvelfya-1* and *fvelfya-2*. Red stars indicate the three critical amino acid sites (312H, 345R, and 387H). B, Relative expression levels of the *FveLFY* homologues in the corresponding lines. One negative control in *lfy-5* was shown; the other three controls without any amplification were omitted. Data are the mean ±SD of three biological replicates. C, Phenotypes of the *lfy-5* and *lfy-5 FveLFYs-ox* plants. Red arrowheads indicate solitary flowers generated from leaf axils.

To test the functional conservation of *FveLFY*s, they were overexpressed in the *Arabidopsis lfy-5* mutant (L*er*-0) under the control of CaMV 35S promoter. The *FveLFY* overexpression lines were selected and validated by qRT-PCR (Figure 3B). In the *FveLFYa-ox* and *FveLFYb-ox* transgenic lines, the primary shoots terminate with a few flowers, and secondary shoots are replaced by solitary flowers (Figure 3C), resembling the *AtLFY-ox* plants (Weigel and Nilsson, 1995). By contrast, neither *FveLFYc* nor *FveLFYd* can rescue the *lfy-5* flower defects. These results suggest that only FveLFYa and FveLFYb are functionally conserved with AtLFY.

### The abundantly expressed *FveLFYc* has no discernable functions

To elucidate the roles of *FveLFYa*–*d*, we analyzed their expression profiles according to a publicly available transcriptome database (Li et al., 2019). *FveLFYa* is highly expressed in floral meristem (FM), receptacle meristem (REM), carpels and SAM (Figure 4A). *FveLFYb* transcripts were only detected in FM and REM at a very low level. *FveLFYc* transcripts were substantially accumulated in FM, REM and carpels. *FveLFYd* transcripts were hardly detectable. The expression levels of *FveLFY*s in floral buds and young leaves were confirmed by qRT-PCR (Figure 4B). To investigate the functions of *FveLFYc*, we performed CRISPR/Cas9-mediated gene editing and generated two *fvelfyc* alleles. The two *fvelfyc* alleles have an A or T insertion at the same site in the first exon (Figure 4C), which cause frameshifts in the N-terminal domain. Therefore, *fvelfyc-1*^*CR*^ and *-2* ^*CR*^ should be strong or null mutants of *FveLFYc*. According to our observation, both *fvelfyc* mutants appear completely normal in the morphology of flowers as well as other organs (Figure 4D). Moreover, the *fvelfya-1 fvelfyc-1*^*CR*^ or *fvelfya-1 fvelfyc-2* ^*CR*^ flowers are indistinguishable from those of *fvelfya-1*. These results suggest that *FveLFYc* is not essential for plant growth and development in *F. vesca*.

**Figure 4.**
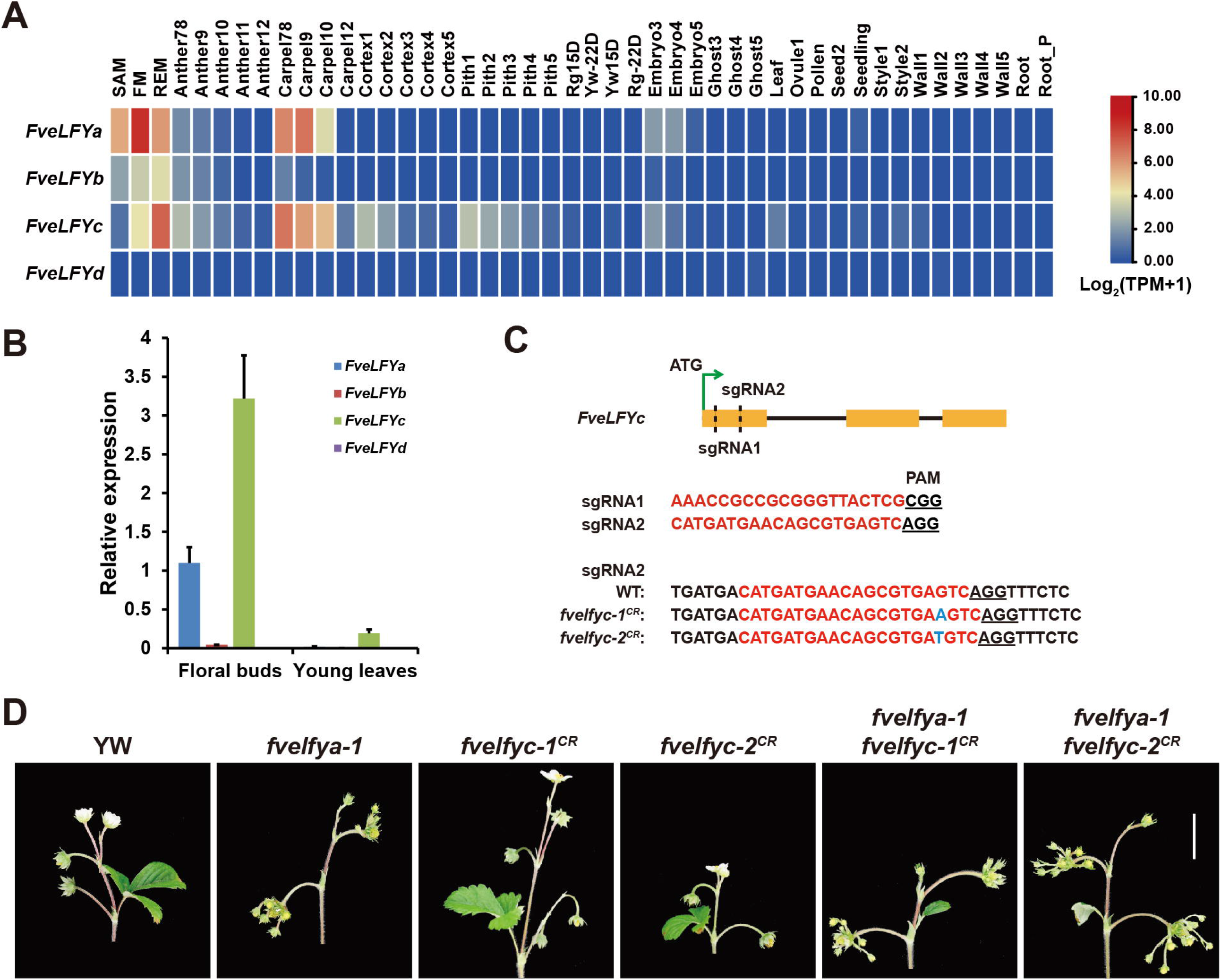
Mutations of *FveLFYc* cause no discernible phenotypes. A, Heat map showing the stage and tissue-specific expression profiles of *FveLFYa-d* according to the transcriptome data from Li et al. (2019). TPM, transcripts per million. B, Relative expression levels of *FveLFYa-d* in floral buds and young leaves of YW. Data are the mean ±SD of three biological replicates. C, CRISPR-Cas9 targeting of *FveLFYc*. Insertions are indicated by blue font. D, Inflorescences of YW, *fvelfya, fvelfyc*, and *fvelfya fvelfyc*. Scale bar: 2 cm.

### Transcription of *FveAP1* is not regulated by *FveLFYa*

To investigate the mechanism underlying the striking *fvelfya* flower defects, we performed transcriptome analysis for the flower buds of *fvelfya-1* and YW at stages 1-6, when the ectopic florets are not formed yet (Supplemental Table S1). From this, we identified 424 significantly downregulated genes and 227 significantly upregulated genes in *fvelfya-1* vs YW (Supplemental Figure S4A and Supplemental Dataset S1). The downregulated genes have enriched GO terms of lignin and other compounds catabolic processes (Supplemental Figure S4B). The upregulated genes have enriched GO terms of defense response and response to different stimuli (Supplemental Figure S4C). Similarly, LFY reduces defense responses in *Arabidopsis* (Winter et al., 2011). This suggests that FveLFYa may play a conserved role in defense response.

Among the differentially expressed genes, the class B flower development genes *AP3a* (FvH4_1g12260), *PIa* (FvH4_2g27860), and *PIb* (FvH4_2g27870) were significantly downregulated in *fvelfya-1* compared to wild type (Figure 5A). In contrast, the expression levels of *AP1* (FvH4_4g29600) and *AG* (FvH4_3g06720) show no statistical difference between *fvelfya-1* and YW. To confirm these, we performed qRT-PCR with floral buds of YW, *fvelfya-1* and *FveLFYa-ox*. Consistent with the RNA-seq results, expression of *FveAP1* shows no difference among YW, *fvelfya-1* and *FveLFYa-ox* lines (Figure 5B). The B class genes (*AP3a, AP3b-*FvH4_2g38970, *PIa* and *PIb*) were significantly downregulated in *fvelfya-1* and slightly upregulated in *FveLFYa-ox* plants, compared to those of YW. However, *FveAG* was significantly downregulated in *fvelfya-1* floral buds in qRT-PCR (Figure 5B), therefore the expression level of this gene in RNA-seq data was somehow inaccurate. These results indicate that wild type *FveLFYa* functions to activate the class B and C gene expression, but loses the capacity to regulate *FveAP1* expression.

**Figure 5.**
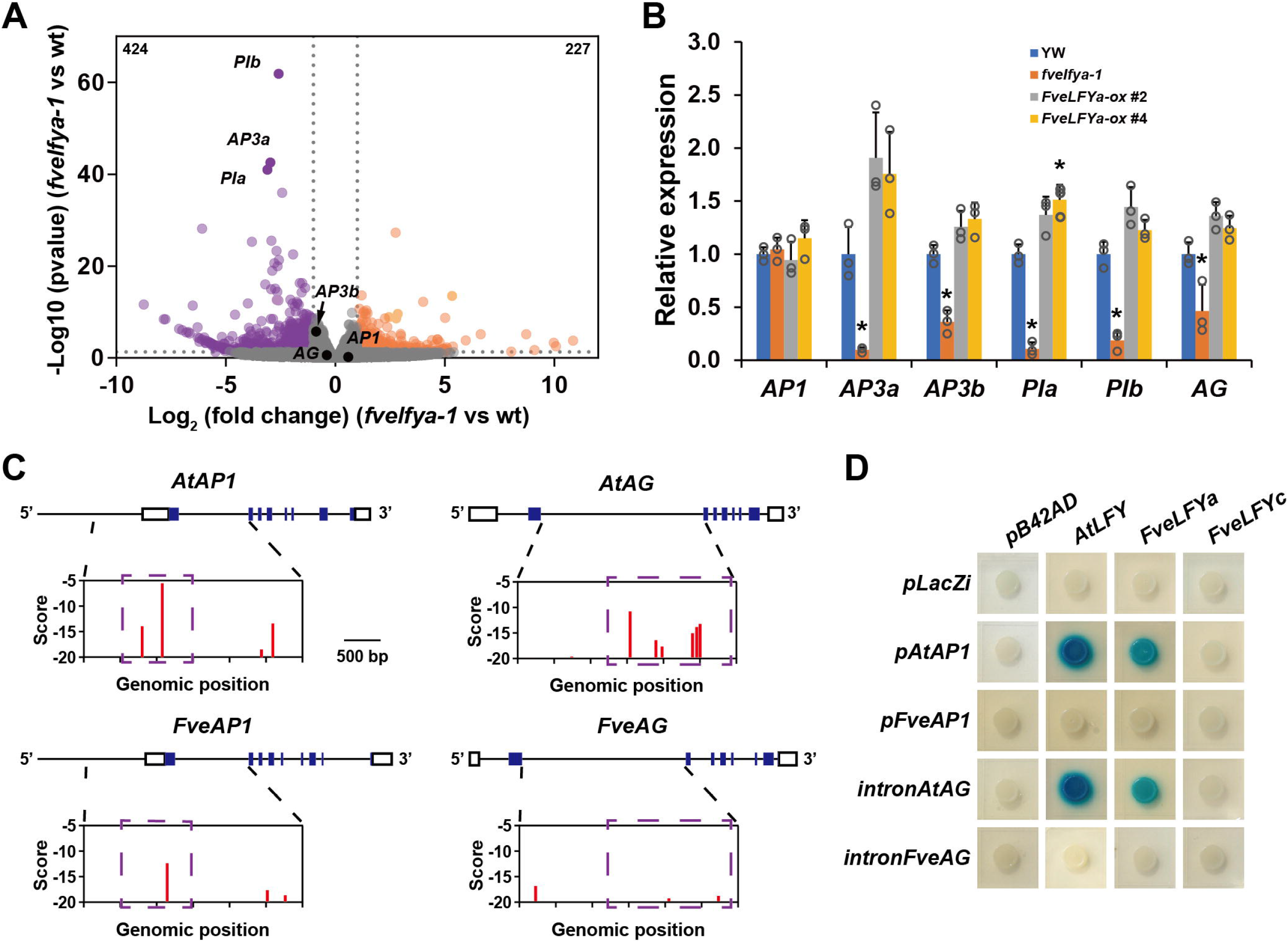
Divergent regulation of *FveAP1* and *FveAG* mediated by *FveLFYa*. A, Volcano plot shows differentially expressed genes in floral buds of *fvelfya-1* relative to YW. Purple dots represent downregulated genes in *fveflya-1*; orange dots represent upregulated genes in *fveflya-1*. The vertical dashed lines mark Log_2_FC of -1 and 1, and the horizontal dashed lines mark an adjusted P value of 0.05. B, Expression levels of *FveAP1, FveAP3a, FveAP3b, FvePIa, FvePIb*, and *FveAG* in floral buds of WT, *fvelfya-1*, and *FveLFYa-ox* examined by qRT-PCR. Data are the mean ±SD of three biological replicates. *, *P* < 0.01, Student’s *t*-test. C, LFY binding sites prediction in the genomic regions of *AP1* and *AG* in *F. vesca* and *Arabidopsis*. Dashed rectangles indicate the key LFY-binding regions. Open boxes indicate noncoding sequences in the transcript, and closed boxes indicate coding sequences. Scores are computed with the SYM-T model. D, Yeast one-hybrid assays showing the binding activities between LFYs and the *AP1* promoters or the large introns of *AG* homologues. The yeast cells were grown on the SD-Trp-Ura plates containing X-gal.

Previous studies revealed that LFY regulates the expression of *AP1* and *AG* through direct binding to the promoter of *AP1* and the large second intron of *AG* in *Arabidopsis* (Moyroud et al., 2011). The LFY/*AP1* and LFY/*AG* transcriptional links have been found to be conserved in several species (Moyroud et al., 2011; Monniaux et al., 2017). Using the SYM-T scoring matrix, we predicted the LFY binding sites in the genic regions of *AtAP1, AtAG, FveAP1*, and *FveAG*. Consequently, the LFY binding sites are shown to be reduced in number and have lower affinity in the promoter and first intron of *FveAP1* and the large intron of *FveAG* when compared to the *cis*-elements of *AtAP1* and *AtAG*, respectively (Figure 5C). To validate the predictions, yeast one-hybrid assay was performed. While both AtLFY and FveLFYa interact with the *AtAP1* promoter and the intron of *AtAG*, neither of them interacts with the corresponding regions of *FveAP1* and *FveAG* (Figure 5D). As negative controls, FveLFYc can bind neither the *AP1* promoters nor the *AG* introns (Figure 5D). In addition, the prediction revealed that there are potential binding sites of FveLFYa in the promoter and 3’ end of *FveAG* (Supplemental Figure S5), consistent with the reduced expression of *FveAG* in *fvelfya* (Figure 5B). Together, *FveAP1* transcription is independent of *FveLFYa* owing to the loss of FveLFYa binding sites in the *FveAP1* promoter, explaining why *FveAP1* expression is unchanged in the *fvelfya* mutants.

### *LAM* is required for ectopic floret formation in *fvelfya*

*LAM* encodes a GRAS transcription factor required for stamen and axillary bud initiation in *F. vesca* (Feng et al., 2021). To explore the potential genetic interaction between *FveLFYa* and *LAM*, the *lam* mutant was crossed with the *fvelfya-2*/+ heterozygote. In the *fvelfya-2 lam* double mutant, one notable change is that the number of sepals or sepal-like organs was largely reduced compared to *fvelfya-2* (Figure 6A). This reduction in sepal number in *fvelfya-2 lam* versus fvelfya-2 corresponded very closely to the number of stamens in a typical wild-type flower (Figure 6B). These results suggest that most of the extra sepals or sepal-like organs in *fvelfya* flowers are converted from stamens. Moreover, the ectopic florets were almost eliminated in *fvelfya-2 lam* (Figure 6A, C), indicating that *LAM* is required for floret initiation at the sepal axils in *fvelfya* flowers. It was reported that *AP1* overexpression contributed to the outgrowth of secondary flowers in the *lfy* mutants in *Arabidopsis* (Liljegren et al., 1999). According to the yeast-two hybrid and split-LUC assays, FveAP1 could directly interact with LAM (Supplemental Figure S6). This protein-protein interaction may contribute to the reiterative outgrowth of ectopic florets in the *fvelfya* mutants.

**Figure 6.**
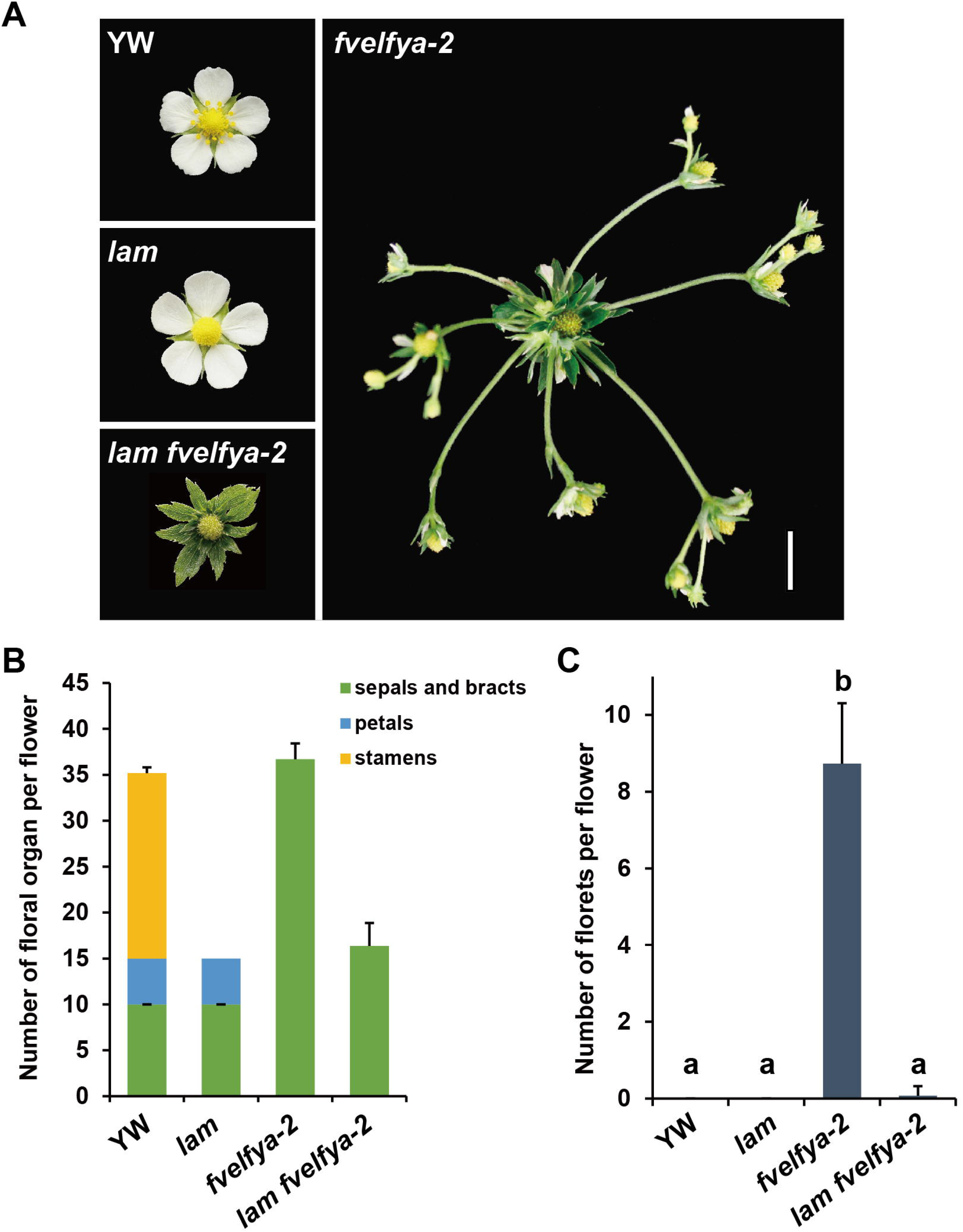
*LAM* is required for the growth of secondary florets in *fvelfya*. A, Flowers of YW, *lam, fvelfya-2* and *fvelfya-2 lam*. Scale Bar: 1 cm. B, Numbers of sepals and bracts, petals, and stamens in each flower of YW, *lam, fvelfya-2* and *fvelfya-2 lam*. n > 15. C, Numbers of ectopic florets in each flower of YW, *lam, fvelfya-2* and *fvelfya-2 lam*. n > 15. Data are the mean ±SD. Different letters indicate significant differences among plant lines (P < 0.01; one-way ANOVA and Tukey HSD).

### *FveLFYa* plays a moderate role in leaflet initiation and expansion in *F. vesca*

Like wild type, most leaves in the *fvelfya* mutants have three leaflets, including one terminal leaflet and two lateral leaflets (Figure 7A). However, abnormal leaves were observed in *fvelfya* after pruning. These leaves can be separated into different types. A majority of abnormal leaves have two smaller lateral leaflets or sometimes one smaller lateral leaflet (Figure 7B). For the severe types, all the three leaflets become much smaller than wild type. In addition, virtually all abnormal leaves have elongated petiolules of terminal leaflets. Next, the percentages of different leaves were recorded in YW and different mutants. While 98% of YW leaves and all the *fvelfyc-1* ^*CR*^ leaves are normal, approximately 30% leaves are abnormal in the three *fvelfya* single mutants and *fvelfya-1 fvelfyc-1*^*CR*^ double mutants (Figure 7C). These results suggest that *FveLFYa*, rather than *FveLFYc*, is required for leaf development in *F. vesca*.

**Figure 7.**
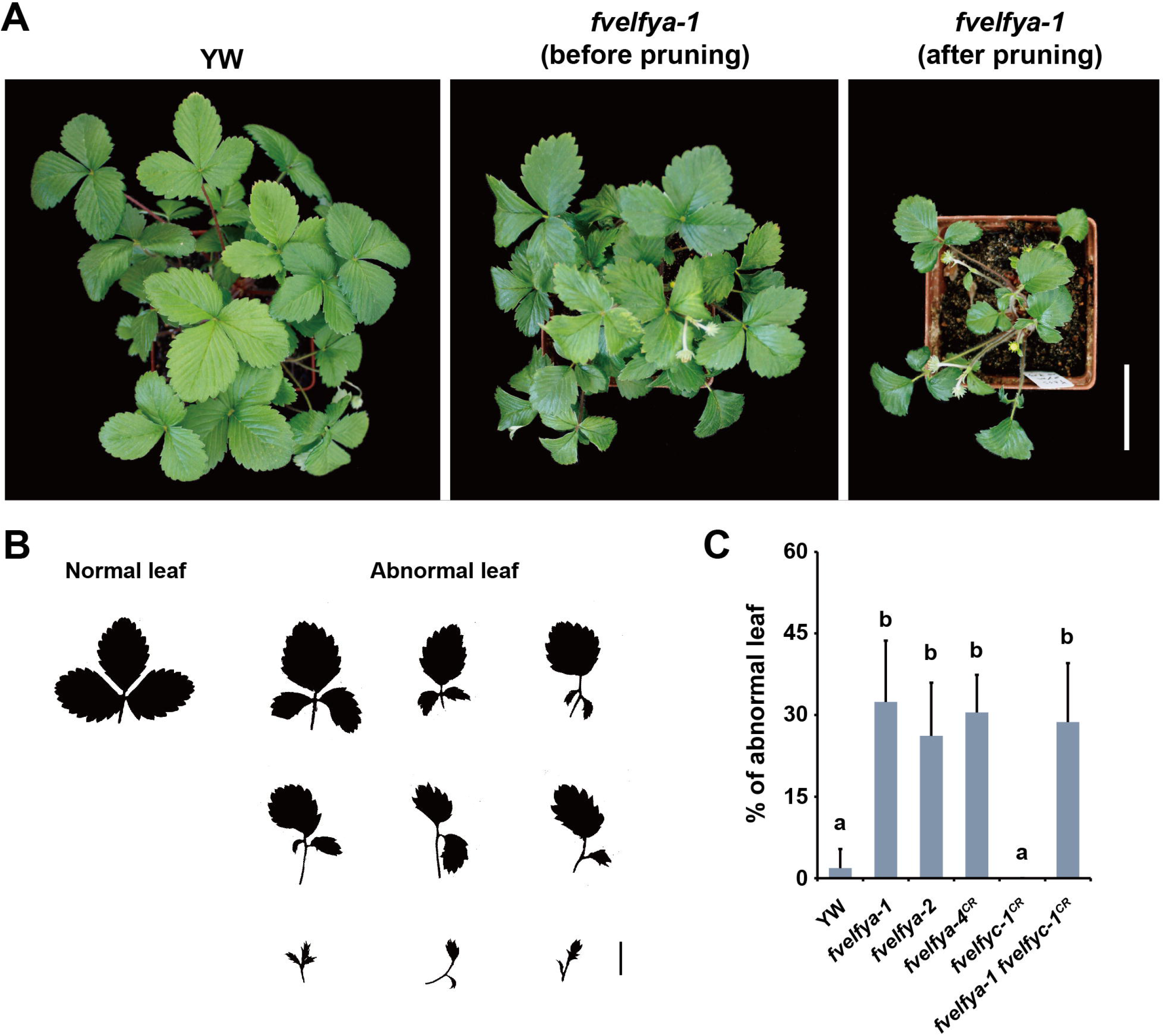
*FveLFYa* regulates leaf development in *F. vesca*. A, Top views of YW and the *fvelfya-1* plant before and after pruning. Scale bar: 5 cm. B, Normal and abnormal leaves in *fvelfya-1*. Scale bar: 2 cm. C, Percentages of abnormal leaves in the indicated genotypes. Data are the mean ±SD. Different letters indicate significant differences among plant lines (P < 0.01; one-way ANOVA and Tukey HSD).

### Evolutionary features of the four *FveLFY* genes

The presence of four *FveLFY* genes in *F. vesca* is unusual, as most species have only a single copy. To determine the phylogeny of these genes, a phylogenetic tree was constructed for a total of 78 *LFY* homologues in angiosperms, mostly from the Rosaceae lineage. In the phylogenetic tree, the *Fragaria LFY* homologues from 7 diploid species were clustered into the *LFY*a, *LFY*b and *LFY*c/d clades (Figure 8A), suggesting that the four *LFY* homologues have arisen before the species divergence in the *Fragaria* genus. By contrast, only a single copy of *LFY* is found in the *Rubus, Rosa* and *Potentilla* genera of Rosoideae subfamily, suggesting that *FveLFY* homologues were duplicated independently within the Potentilleae lineage.

**Figure 8.**
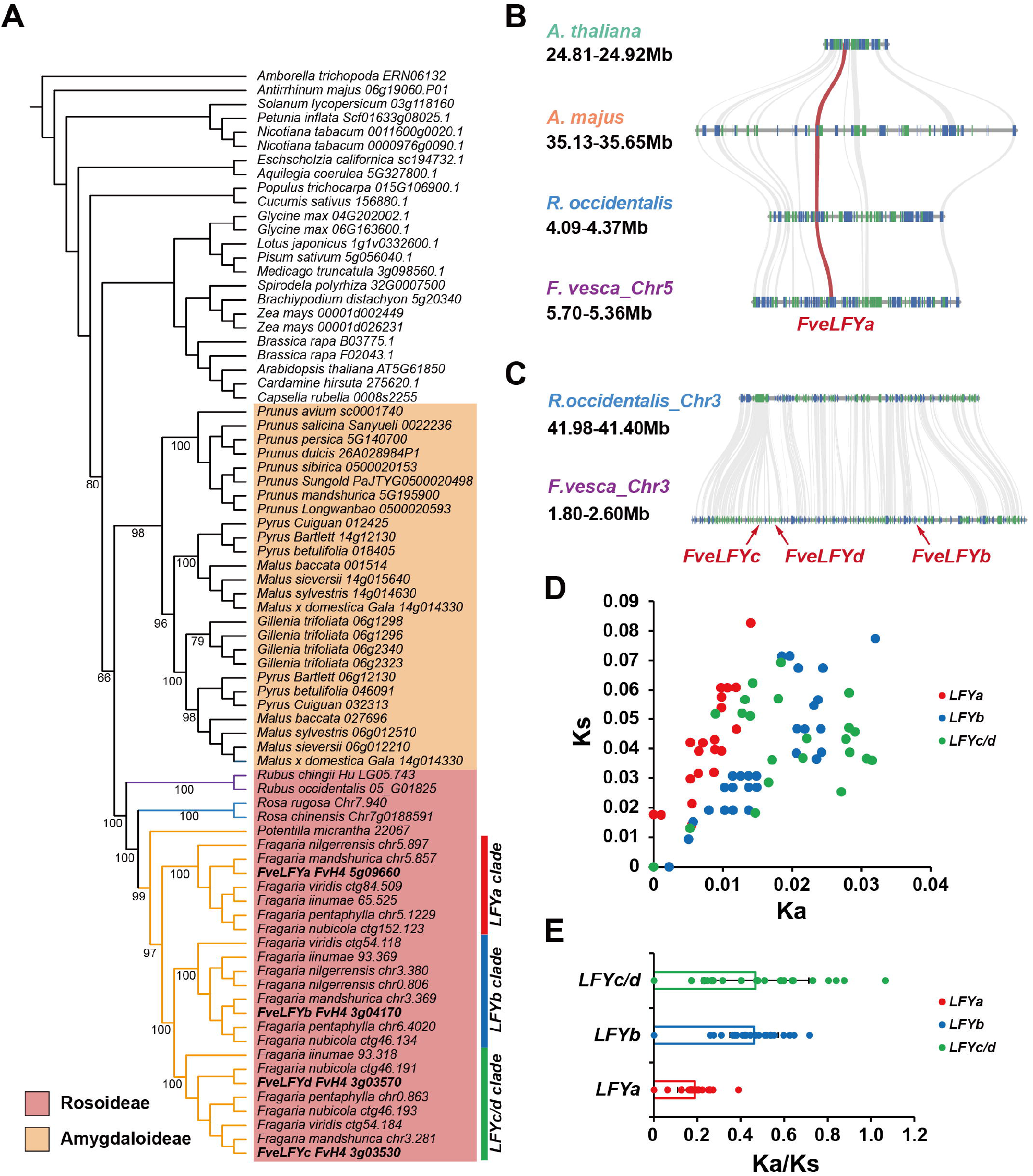
Evolution features of *FveLFYa-d*. A, *LFY* phylogeny within selected angiosperms. Numbers at nodes indicate ML bootstrap values. Purple branches represent the tribe Rubeae, light blue branches represent the tribe Roseae, and orange branches represent the tribe Potentilleae. B, Multiple genomic synteny alignments of the *LFY*-associated genomic blocks in the four selected species. Red strips highlighted the syntenic connections of the *LFY* gene loci. C, Neighboring genomic region of the *FveLFYb*-*d* locus in *F. vesca* is syntenic to the *R*.*occidentalis* genomic region without *LFY*. D, Pairwise Ka and Ks divergence of *LFYa, LFYb* and *LFYc*/*d* clades in *Fragaria*. E, Pairwise Ks/Ks ratio divergence of *LFYa, LFYb* and *LFYc*/*d* clades in *Fragaria*.

Next, synteny analysis was performed for the flanking genomic regions around the *LFY* loci in *A. thaliana, Antirrhinum majus, Rubus occidentalis* and *F. vesca*. The adjacent genomic region of *FveLFYa* was syntenic to the *LFY* genomic regions in the other three species (Figure 8B), suggesting that *FveLFYa* is the *F. vesca LFY* orthologue. By contrast, no synteny was observed in the *FveLFYb-d* adjacent regions between *F. vesca* and *R. occidentalis* (Figure 8C), indicating an independent origin for the three *FveLFY* genes.

To gain insight on *LFY* evolution in *Fragaria* species, we calculated non-synonymous (Ka) and synonymous (Ks) substitution rates and Ka/Ks ratios for gene pairs in the *LFYa, LFYb, LFYc*/*d* clades, respectively. The Ks values of the *LFYa, LFYb* and *LFYc*/*d* clades were located in a similar range (Figure 8D, Supplemental Dataset S2). By contrast, the Ka values of the *LFYb* and *LFYc*/*d* clades were higher than those of the *LFYa* clade. Furthermore, the average Ka/Ks ratio of *LFYa* gene pairs was 0.19, while those of *LFYb* and *LFYc*/*d* gene pairs were 0.46 and 0.47 (Figure 8E). These results indicate that LFYa clade genes underwent strong purifying selection, while *LFYb* and *LFYc*/*d* genes underwent relaxed constraints or positive selection.

## Discussion

LFY is a well conserved plant-specific transcription factor and a key regulator of flowering and floral patterning. Recent studies showed that LFY acts as a pioneer transcription factor to access its binding sites even in the closed chromatin regions and recruit chromatin remodelers (Jin et al., 2021; Lai et al., 2021), highlighting the importance of this gene. In most cases, *LFY* is a single copy gene in each genome. The woodland strawberry genome however possesses four *LFY* homologues, *FveLFYa-d*, offering a rare opportunity to investigate the functional conservation and evolution of LFY in flowering plants. In this study, a combination of genetic and molecular approaches was used to demonstrate that *FveLFYa* plays a predominant role in floral patterning and leaf development in woodland strawberry.

### Disruption of *FveLFYa*, rather than other *FveLFY*s, causes special floral morphology

*FveLFYa* is the *LFY* orthologue in *F. vesca* according to the results of amino acid alignment, microsynteny analysis, and ectopic expression in *Arabidopsis*. The *fvelfya* flowers show homeotic conversion of floral organs and massive outgrowth of reiteratively formed ectopic florets in sepal axils (Figure 1). In comparison, most reported mutants of the *LFY* orthologs in a variety of flowering species show partial homeotic conversions of flowers to shoots, and commonly have secondary flowers in sepal axils (Weigel et al., 1992; Hofer et al., 1997; Dong et al., 2005; Wang et al., 2008; Ikeda-Kawakatsu et al., 2012; Zhao et al., 2016). Nevertheless, transformation of carpels to bract-like organs and massive outgrowth of ectopic florets are special phenotypes to strawberry.

In the gene network of *FveLFYa*, downregulation of the B class genes *AP3a, AP3b, PIa*, and *PIb* in *fvelfya* is consistent with the homeotic conversion of petals and stamens. The C class gene *FveAG* was also significantly inhibited in *fvelfya*, which might be owing to the presence of LFY binding sites in the promoter or 3’-end (Supplemental Figure S5), rather than in the second intron (Figure 5C, D). These results indicate that activation of the B and C class genes by LFY is well conserved in *F. vesca*. LFY and the interacting co-factor UFO work together to activate *AP3* for petal and stamen identity specification (Lee et al., 1997; Chae et al., 2008). The B and C genes were not significantly up-regulated in the *FveLFYa-ox* transgenic lines (Figure 5B), indicating that other factors might be required, such as *UFO*. Similar to the *fvelfya* flowers, the *fveufo1* flowers show partial homeotic conversion of floral organs, elongated internodes and ectopic flower (Shahan et al., 2018). Therefore, the joint roles of LFY and UFO in floral patterning probably are also conserved in *F. vesca*.

According to the phylogenetic relationships, the *Fragaria* species can be divided into north and south clade (Qiao et al., 2021; Sun et al., 2021). Both *F. nubicola* in the south clade and *F. vesca* in the north clade contain the LFYa-d homologues, thus we propose that they had duplicated in the common *Fragaria* ancestor. Among the four *F. vesca* LFY homologues, FveLFYa and FveLFYb belong to the type I LFY with similar functions to AtLFY based on the protein sequence analysis and ectopic expression in *Arabidopsis*. In spite of this, *FveLFYb* seems unlikely to play important roles in strawberry growth and development due to its very low expression levels. Likewise, FveLFYc and FveLFYd may have minimal or no function, as they failed to rescue *Arabidopsis lfy-5* mutant (Figure 3). Moreover, the *fvelfyc*^*CR*^ mutants look completely normal (Figure 4D). The reason might be that FveLFYc is not a type I LFY based on the amino acid sequence, and has lost the conserved DNA binding ability for *AtAP1* and *FveAP1* (Figure 5D). *FveLFYd* is a pseudogene and barely expressed according to the published transcriptomic data (Li et al., 2019).

### *FveLFYa* can’t regulate *AP1* expression in *F. vesca*

In *Arabidopsis*, LFY directly activates *AP1* expression through binding to the promoter region (Parcy et al., 1998; Wagner et al., 1999; Moyroud et al., 2011; Winter et al., 2011). Unexpectedly, in *F. vesca, FveAP1* expression was not altered in *fvelfya* compared to WT. Our LFY binding site predictions and yeast one hybrid assays indicate that changes in the *FveAP1* promoter sequence contribute to the inability of FveLFYa to bind and regulate *FveAP1* expression. Similarly, changes in the large intron of *FveAG* have contributed to the inability of FveLFYa to bind *AG* intron (Figure 5), which contrasts with evolutionarily conserved *AG* intron elements for LFY binding in other flowering species (Moyroud et al., 2011). These findings suggest that changes in the *cis*-regulatory sequences of target genes might be an important mechanism for the changes of regulatory network underlying variation of floral organ development.

According to previous studies, expression of *FveAP1* in *fvelfya* may underlie the reiterative outgrowth of ectopic flowers. In *Arabidopsis*, the *lfy* mutants showed a partial homeotic conversion of flower organs and formation of secondary flower buds in sepal axils, while the *lfy ap1* double mutants showed a complete conversion from floral organs to leaves at the same position (Schultz and Haughn, 1991; Weigel et al., 1992). We speculate that the secondary flowers formed in *lfy* are caused by the residual expression of *AP1*. Conversely, constitutive expression of *AP1* in *lfy* induced massive outgrowth of secondary flowers (Liljegren et al., 1999). In *C. hirsuta*, mutations in *ChLFY* cause a flower to shoot conversion, where expression of *AP1* is completely eliminated (Monniaux et al., 2017). These results suggested that the *AP1* dosage is closely associated with the flower development defects in *lfy*. These fit well to “the rate of meristem maturation” model, which proposed that meristem maturation is a gradual process (Park et al., 2012). According to the model, we hypothesize that loss of *LFY* delays the meristem maturation, which results in an extended period of indeterminacy, and expression of *AP1* therefore contribute for initiation of ectopic flowers, and *vice versa*.

Mutation of LAM, encoding a GRAS transcription factor, resulted in lack of stamens in *F. vesca* (Feng et al., 2021). Thus, the reduced number of sepals in *fvelfya lam* double mutant relative to *fvelfya* is likely due to the loss of the stamen whorl caused by *LAM* mutation. LAM and its orthologues in *Arabidopsis*, tomato, and rice are the key positive regulators of axillary meristem initiation (Schumacher et al., 1999; Greb et al., 2003; Li et al., 2003). Formation of the ectopic florets was almost suppressed in the *fvelfya-2 lam* flowers (Figure 6), indicating that the *fvelfya* flowers might be partially converted into shoots. As LAM physically interacts with FveAP1 (Supplemental Figure S6), we speculate that their protein complex may mediate the secondary flower formation in sepal axils, and thus the *fveap1 fvelfya* double mutant might have similar phenotypes to *lam fvelfya* in flower development.

### *FveLFYa* regulates leaf development in *F. vesca*

The essential roles of *LFY* homologues in leaf development come from the studies of *UNI* in *P. sativum* and *SGL1* in *M. truncatula*, which are able to determine the formation of compound or simple leaves (Hofer et al., 1997; Wang et al., 2008). In addition, *LFY* has been found to play a role in maintaining leaf complexity in distant angiosperms species, including Fabaceae (Hofer et al., 1997; Wang et al., 2008), Solanaceae (Molinero-Rosales et al., 1999), and Brassicaceae (Monniaux et al., 2017). Like *M. truncatula* leaves, the strawberry leaves are compound with three leaflets. We observed that approximately 30% of mature leaves in each *fvelfya* plant are abnormal with fewer and smaller leaflets (Figure 7), suggesting that *LFY* also takes part in leaf development in the Rosaceae family. The weak leaf phenotypes of *fvelfya* suggest that *FveLFYa* either plays a moderate role or has more impacts but covered by other genes in this process. Nevertheless, this result leads us to hypothesize that the role of *LFY* in leaf development is ancestral.

In summary, our study makes a few new findings on the *LFY* homologues in strawberry. The *fvelfya* mutants exhibit a special flower phenotype showing massive outgrowth of ectopic florets. Among the four *LFY* homologues in *F. vesca*, only *FveLFYa* has discernable functions. The iterative formation of ectopic florets in *fvelfya* might be related to the constitutive expression of *FveAP1* and require the GRAS transcription factor LAM. *FveLFYa* also takes part in leaf development. The *FveLFYa* homologues might have duplicated prior to the divergence of the extant *Fragaria* species, but stay nonfunctional owing to low expression levels, amino acid substitutions, or gene structural changes. These results deepen our understanding about the functions and evolution of this important low-copy transcription factor.

## Materials and Methods

### Plant material and growth conditions

The inbred lines of the *F. vesca* varieties Yellow Wonder (YW) and Hawaii-4 (H4) were used as wild type. The *lam* mutant in *F. vesca* was previously described (Feng et al., 2021). The *Arabidopsis lfy-5* mutant in Landsberg *erecta* (L*er*) background was obtained from Prof. Xiaolan Zhang (Weigel et al., 1992). Seeds were stratified in water for two days for *Arabidopiss* or 7-10 days for *F. vesca* at 4°C and then sowed on the soil. Plants were grown under long-day conditions (16 h light; 8 h darkness) with a light intensity of 100 μmol m^-2^ s^-1^ at 22°C.

### Paraffin sections

Floral buds were infiltrated with FAA solution (50% ethanol, 5% acetic acid, 10% formaldehyde) under vacuum for 30 min and incubated for 24 h. The samples were sequentially dehydrated through a series of ethanol solutions (30%, 50%, 70%, 85%, 95%, and 100%), cleared with xylene and embedded in paraffin. Sections were obtained using a Leica Microtome, de-waxed in xylene, rehydrated, stained with 0.5% toluidine blue for 10-20 s and then sealed with resin. Images were taken using a Zeiss Axio Scope A1 microscope.

### Gene isolation of Y535 and Y904 mutants

Y535 and Y904 were backcrossed to the wild type variety YW, respectively. In the F_2_ populations, the mutants were identified as flowers with extra sepals converted from petals and stamens and ectopic florets. Equal amounts of young leaves were pooled from 20 F_2_ Y535 mutants, 15 F_2_ Y904 mutants, and their own wild-type like segregants (∼20 plants) respectively to extract the genomic DNA using a CTAB method. The candidate SNPs for Y535 and Y904 were identified as previously described (Feng et al., 2021) and further confirmed by Sanger sequencing in more F_2_ mutant-looking individuals.

### Plasmid construction and plant transformation

For *35S::FveLFYa* transformed into strawberry, the genomic fragment (2125 bp) of *FveLFYa* was amplified by PCR and inserted into the binary vector pK7WG2D. For CRISPR/Cas9 gene editing constructions, sgRNAs for targeting *FveLFYa* and *FveLFYc* were designed using the web server CRISPR-P2.0 (Lei et al., 2014). Pairs of oligonucleotides including targeting sequences were synthesized as primers (Supplementary Table1). Fragments were amplified by PCR using pCBC-DT1DT2 as the template and then inserted into *Bsa*I-digested pKSE401G by the Golden Gate assembly (Tang et al., 2018). For *FveLFY*s overexpression in *Arabidopsis*, the genomic region of each *FveLFY* homologue without stop codon was amplified and inserted into the binary vector pRI101. All constructs were transformed into *Agrobacterium tumefaciens* strain GV3101. *Arabidopsis* plants were transformed by floral dip method (Clough and Bent, 1998). *F. vesca* plants were transformed as previously described (Feng et al., 2018).

### Measurement of leaf size

After all the individuals started to flower, mature leaves were harvested and laid flat on A4 papers with transparent tape. The papers with leaves were scanned (EPSON Perfection V6000) and measured by ImageJ (https://imagej.nih.gov/ij/).

### RNA extraction and qRT-PCR

RNA was extracted using HiPure Plant RNA Mini Kit (Magen). Genomic DNA removal and reverse transcription were performed with PrimeScript TM RT regent kit (Takara). qPCR reaction was performed using iTaq Universal SYBR Green Supermix (Bio-Rad) on a Roche LightCycler 480. FvH4_4g24420 encoding *GAPDH* in *F. vesca* (Lin-Wang et al., 2014) and AT3G18780 encoding *ACTIN2* in *A. thaliana* were used as internal reference genes. Oligonucleotide primers used are listed in Supplementary Table S2.

### RNA-seq and data analysis

Floral buds at flower stages 1-6 of YW and Y535 were sampled, respectively. Approximately 100 floral buds were pooled per sample. Three biological replicates were collected per genotype. Total RNA was extracted using HiPure Plant RNA Mini Kit (Magen). TruSeq RNA libraries were generated and sequenced by Personal Biotechnology Co. (Shanghai, China) using the Illumina NovaSeq platform. Raw reads were processed using Trimmomatic (Bolger et al., 2014) to remove adapter sequences and then mapped against the *F. vesca* reference genome v4.0.a2 (Li et al., 2019) using the program HISAT2 (Kim et al., 2015). Raw reads were counted by featureCounts (Liao et al., 2014) and normalized to transcripts per million (TPM) values using TBtools (Chen et al., 2020). Differential gene expression (DEG) analysis was performed with DEseq2 (FDR < 0.05, fold change > 2) (Love et al., 2014). TBtools was used for heatmaps drawing and GO enrichment analysis (Chen et al., 2020).

### LFY binding site prediction

Genomic regions comprising the 3 kb upstream of the translational start sites, gene bodies, and 3 kb downstream of the stop codon were extracted from *Arabidopsis* and *F. vesca* genomes using TBtools (Chen et al., 2020). For each putative LFY target gene, all motifs matched to the 19-bp LFY binding consensus sequence NNNNNNHBDNHRDNNNNNN were extracted with Geneious Prime (https://www.geneious.com/). Scores of the identified motifs were calculated based on the SYM-T scoring matrix using Perl (Moyroud et al., 2011).

### Yeast one-hybrid assay

The full-length coding sequences of *AtLFY, FveLFYa* and *FveLFYc* were amplified and cloned into the pB42AD vector by homology recombination. The promoter regions of *AtAP1* (−899 to -20 bp) and *FveAP1* (−927 to -16 bp), and the large introns of *AtAG* (+1508 to +3108 bp) and *FveAG* (+1452 to +3147 bp) were amplified and cloned into pLacZi. According to the Yeast Protocols Handbook (Clontech), the activation domain (AD) fusion plasmid and reporter plasmid were cotransformed into the yeast strain EGY48. Transformants were selected on synthetic defined (SD)-Trp-Ura dropout plates, and then transferred onto the plates containing X-gal for transcription activity determination.

### Yeast two-hybrid assay

The coding sequences of *LAM* and *FveAP1* were amplified and cloned into the binding domain (BD) bait vector (pGBKT7) and the AD prey vector pGADT7, respectively. The AD and BD plasmids were cotransformed into yeast strain AH109. Transformants were selected on SD-Leu-Trp plates and then transferred onto SD-Leu-Trp-His-Ade for protein interaction examination.

### Split-Luciferase assay

The *LAM* coding sequence was cloned to the N-terminal luciferase (LUC) fusion vector JW771. The *FveAP1* coding sequence was cloned to the C-terminal LUC fusion vector JW772. The constructs were separately transformed into *Agrobacterium tumefaciens* strain GV3101. The *A. tumefaciens* transformants were co-infiltrated into *N. benthamiana* leaves. Three days later, the infiltrated leaves were sprayed with 1 mM D-Luciferin and dark treated for 10 min. Then the signal was checked by a CCD camera (LB985 NightSHADE).

### Phylogenetic analyses, synteny comparison, synonymous and nonsynonymous divergence estimates

The *LFY* homologues of Rosaceae were collected from the Genome Database for Rosaceae (Jung et al., 2018) using blastn searches and filtered with an E-value threshold of 1e-5. Sequences of other *LFY* homologues were retrieved from Phytozome (https://phytozome-next.jgi.doe.gov/) (Goodstein et al., 2011), LOTUS Base (https://lotus.au.dk/), (http://chi.mpipz.mpg.de/assembly.html) (Gan et al., 2016), and Snapdragon Genome Database (http://bioinfo.sibs.ac.cn/Am/index.php). The *LFY* loci-associated chromosomal regions were selected to generate the genomic synteny plots using the Python JCVI utilities (https://github.com/tanghaibao/jcvi).

The sequences were trimmed and aligned to the existing alignment from Sayou et al. (2014) using MUSCLE (Edgar, 2004). Phylogenetic analyses were conducted with the GTR+I+G substitution model and maximum likelihood from 1000 bootstrap replicates using IQTREE (Nguyen et al., 2014). The data were visualized and edited using ITOL (Letunic and Bork, 2016). Multiple sequence alignment within *LFY*a, *LFY*b and *LFY*c/d were performed with TranslatorX (Abascal et al., 2010) in combination with MUSCLE (Edgar, 2004), respectively. Pairwise Ka, Ks and Ka/Ks values were calculated using KaKs_Calculator 3.0 (Zhang, 2022).

### Data availability

The raw RNA-Seq data in this paper have been deposited in the Sequence Read Archive at NCBI under accession number PRJNA869489. Additional data and biological materials related to this study are available from the corresponding author upon request.

### Statistical analyses

The Student’s *t*-test (two-tailed) was performed between two groups and the one-way ANOVA, Tukey’s post hoc analysis was used for multiple comparisons. These tests were performed using SPSS v22.0 (IBM Crop., Armonk, NY, USA).

## Acknowledgements

We thank Prof. Xiaolan Zhang (China Agricultural University) for providing *Arabidopsis lfy-5* seeds and Prof. Michael Lenhard (Universität Potsdam) for helpful comments on the manuscript.

## Supplemental data

Supplemental Figure S1. Schematic diagram showing the complementation test between Y904 and Y535.

Supplemental Figure S2. Sequence conservation of FveLFYa in the region containing the mutation in Y904.

Supplemental Figure S3. Gene models of *FveLFYa–d*.

Supplemental Figure S4. Differentially expressed genes in the flower buds of *fvelfya-1* compared to the wild type YW.

Supplemental Figure S5. LFY binding sites prediction in upstream and downstream regions of *AtAG* and *FveAG*.

Supplemental Figure S6. FveAP1 directly interacts with LAM.

Supplemental Table S1. Summary of RNA-seq read statistics.

Supplemental Table S2. Primers used in this study.

Supplemental Dataset S1. Differentially expressed genes in the pairwise comparison between fvelfya-1 and YW in flower buds at stages 1-6 (fold change > 2.0, padj < 0.05).

Supplemental Dataset S2. Pairwise synonymous and non-synonymous substitution rates of LFY genes.

## Notes

This work was supported by National Natural Science Foundation of China (32172539), the Fundamental Research Funds for the central Universities (2662022YLPY002), and the China Postdoctoral Science Foundation (2020M682442).

### Competing Interest Statement

The authors have declared no competing interest.

