## Supplemental Figures and Tables for "The roles and evolution of the four *LEAFY* homologues in floral patterning and leaf development in woodland strawberry"

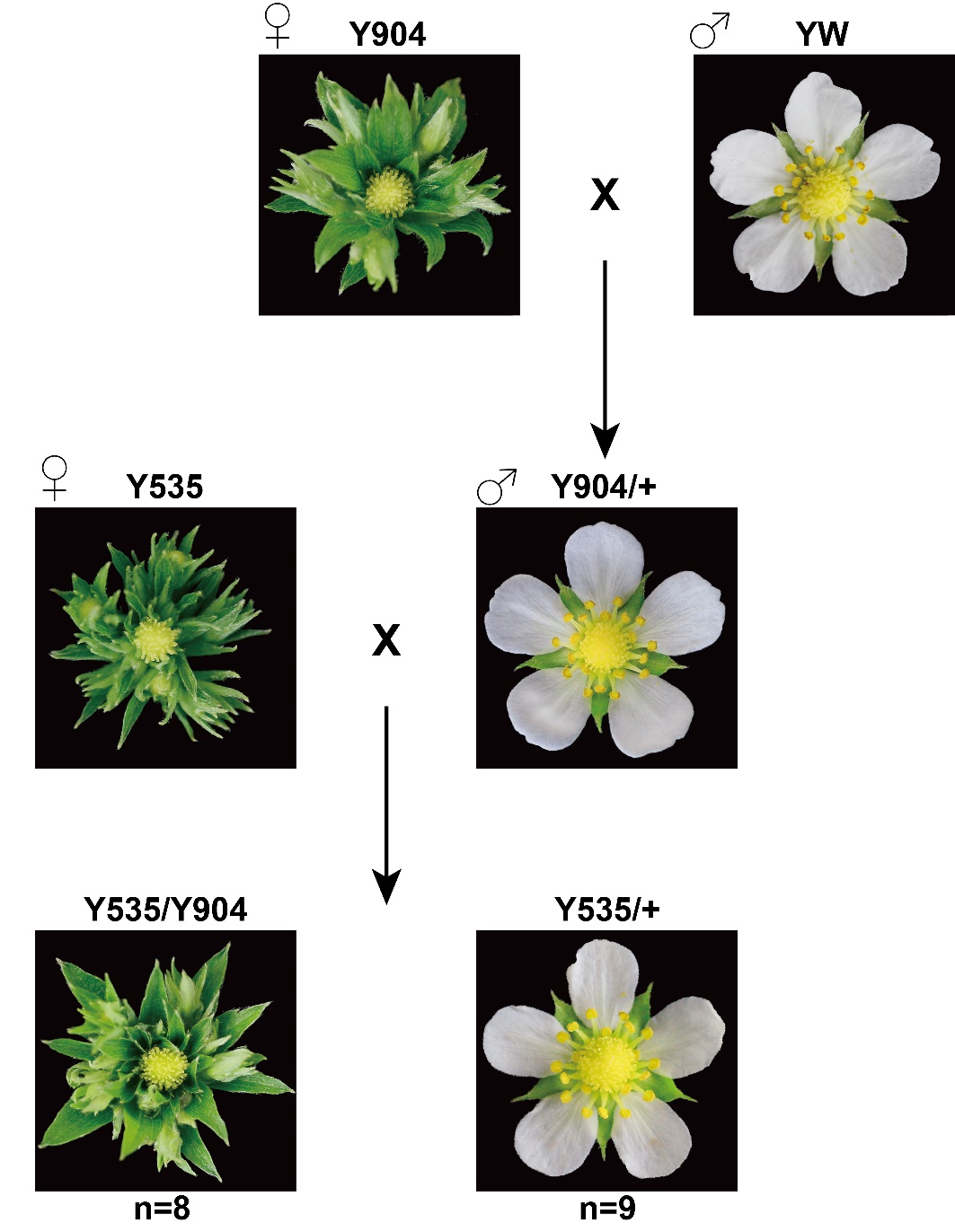

**Supplemental Figure S1. Schematic diagram showing the complementation test between Y904 and Y535.**

Y535 was used as the maternal parent and crossed with the Y904/+ heterozygote. In the F_1_ progeny, eight plants were mutants, and nine plants were wild type, a ratio close to 1:1. The representative flowers for each genotype are shown.

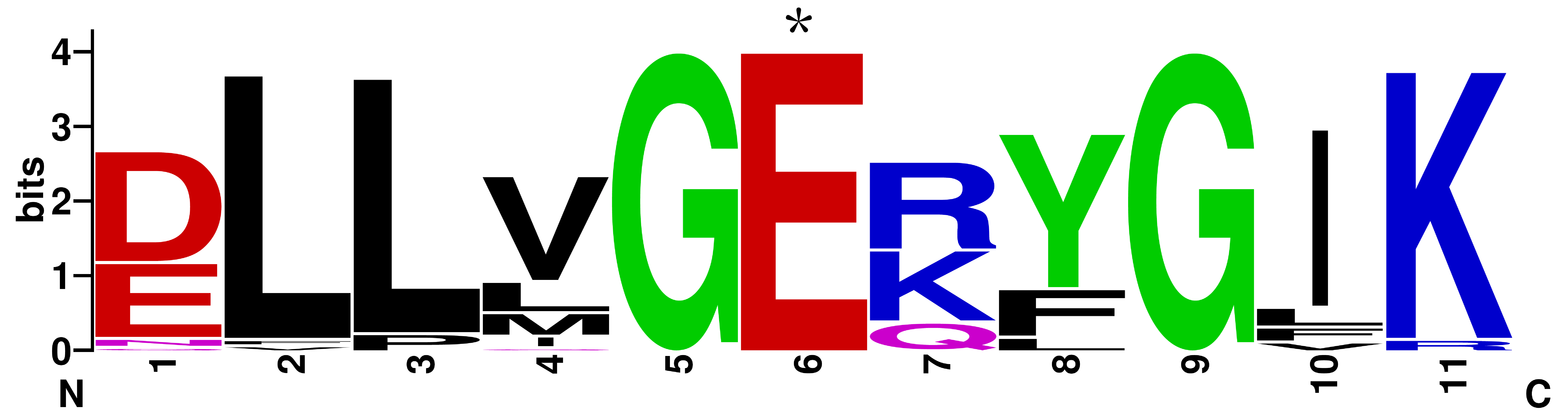

**Supplemental Figure S2. Sequence conservation of FveLFYa in the region containing the mutation in Y904.**

The sequence logo was generated by Weblogo with 40 LFY homologues from algae, hornwort, moss, liverwort, fern, gymnosperm and angiosperm. The asterisk indicates the highly conserved amino acid mutated in Y904.

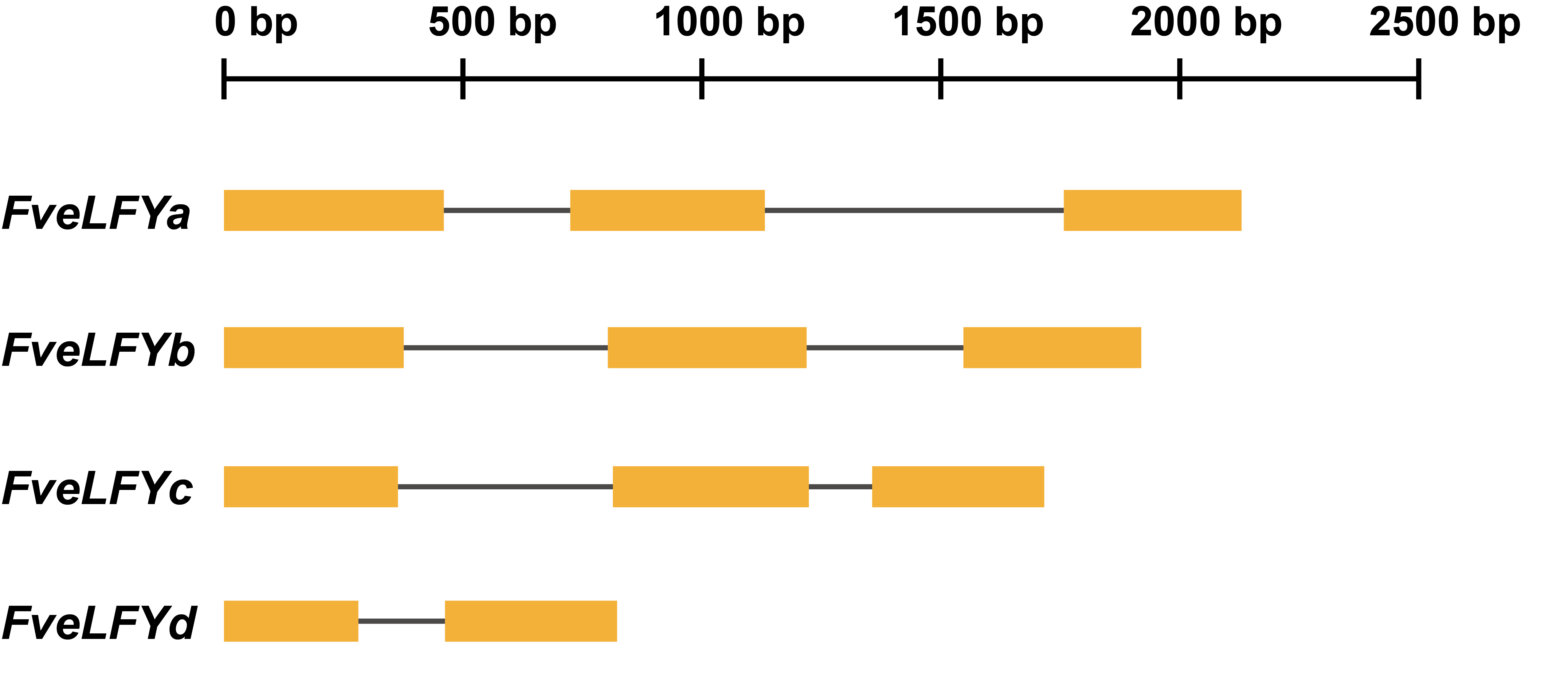

**Supplemental Figure S3. Gene models of *FveLFYa–d*.**

Gene models of *FveLFYa–d*, from the translation start codon to stop codon, were plotted. Yellow boxes indicate the exons, and grey lines indicate the introns.

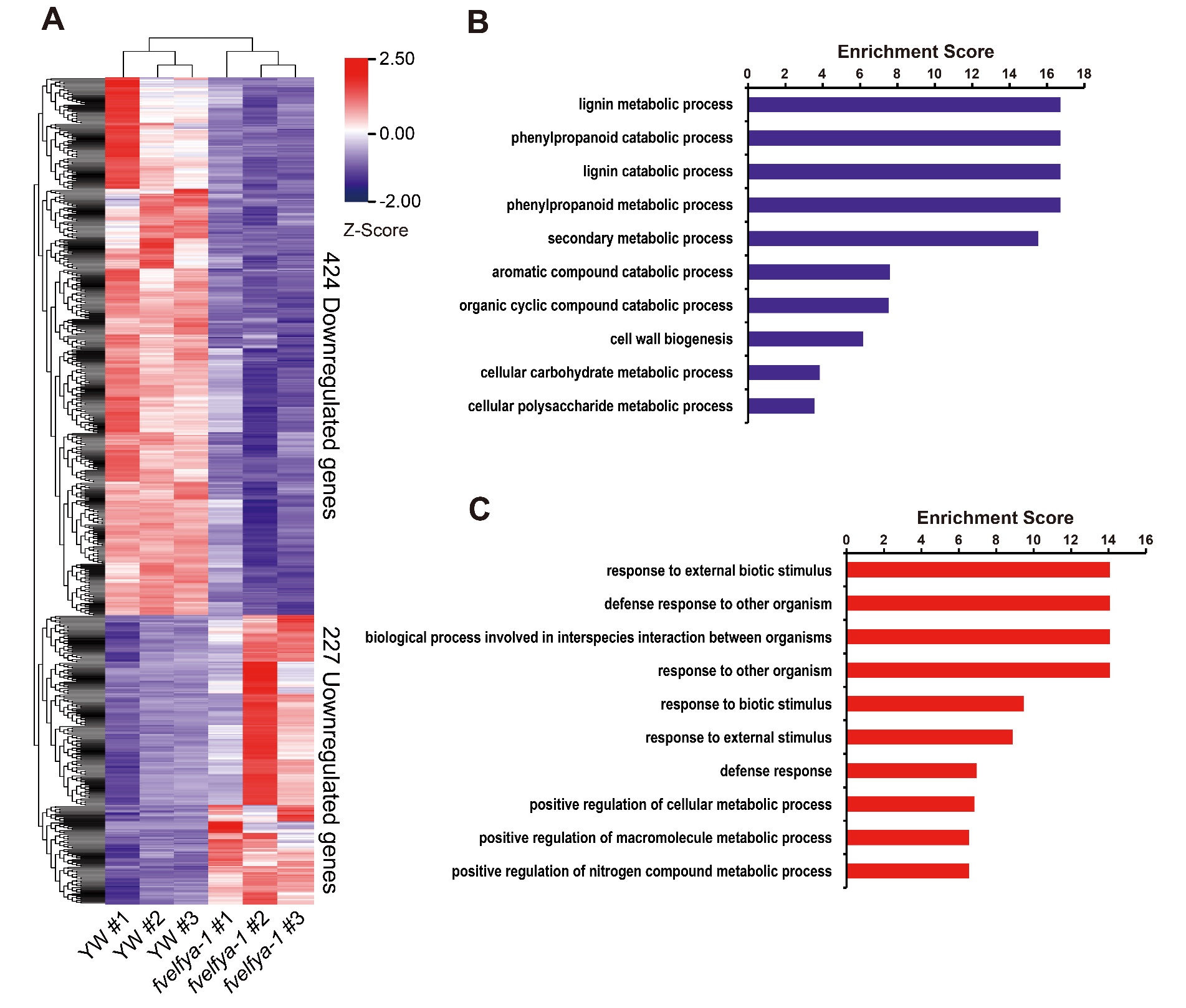

**Supplemental Figure S4. Differentially expressed genes in the flower buds of *fvelfya-1* compared to the wild type YW.**

A, Heatmap showing the expression of 651 differentially expressed genes obtained from RNA-seq with fold change >2 and FDR <0.05. The colors correspond to the Z-Score values calculated from TPM. B and C, Top 10 enriched GO terms in the category of biological process for genes decreased (B) and increased (C) in *fvelfya-1*.

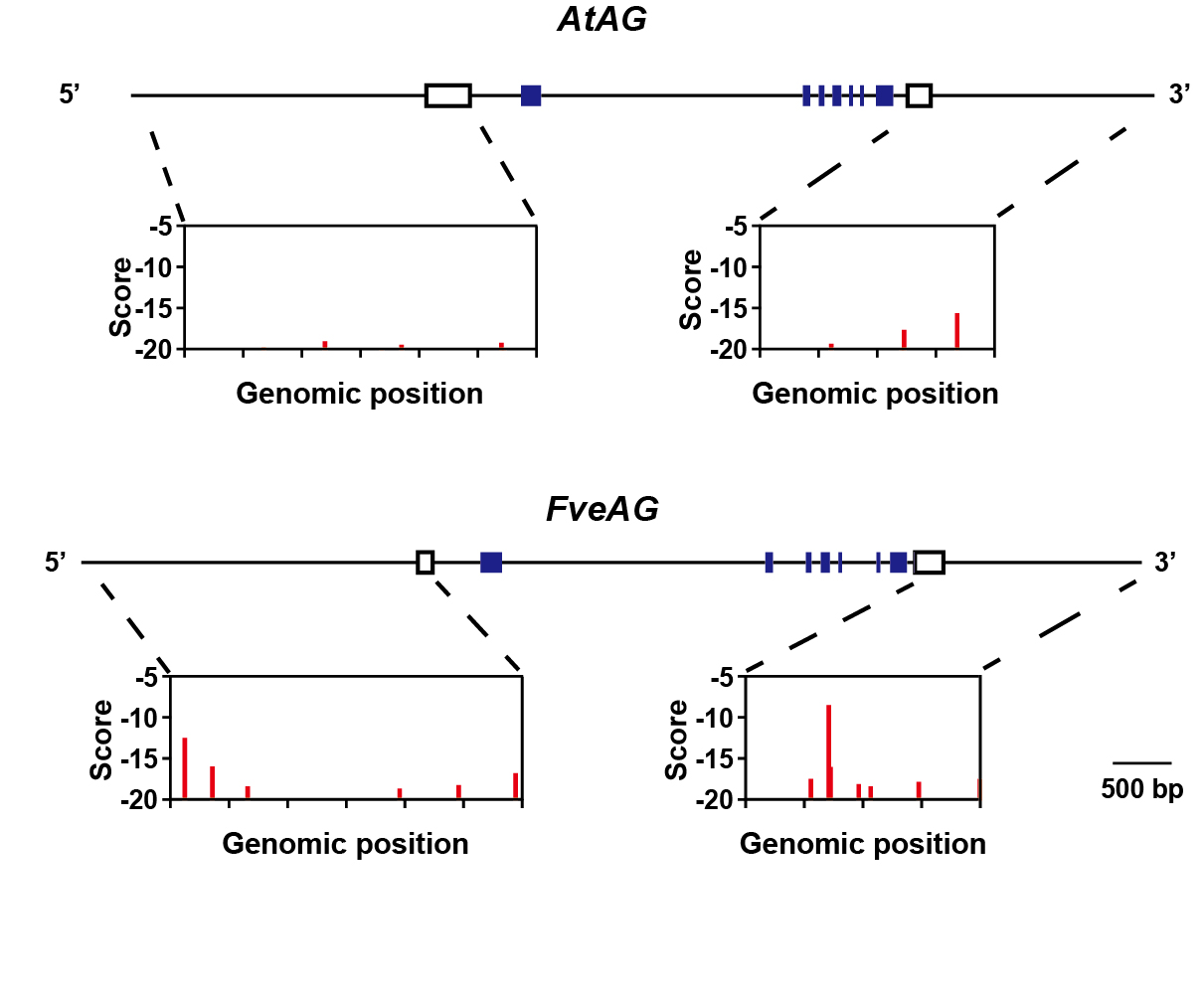

**Supplemental Figure S5. LFY binding sites prediction in upstream and downstream regions of *AtAG* and *FveAG*.**

Open boxes indicate noncoding sequences in the transcript, and closed boxes indicate coding sequences. Scores are computed with the SYM-T model.

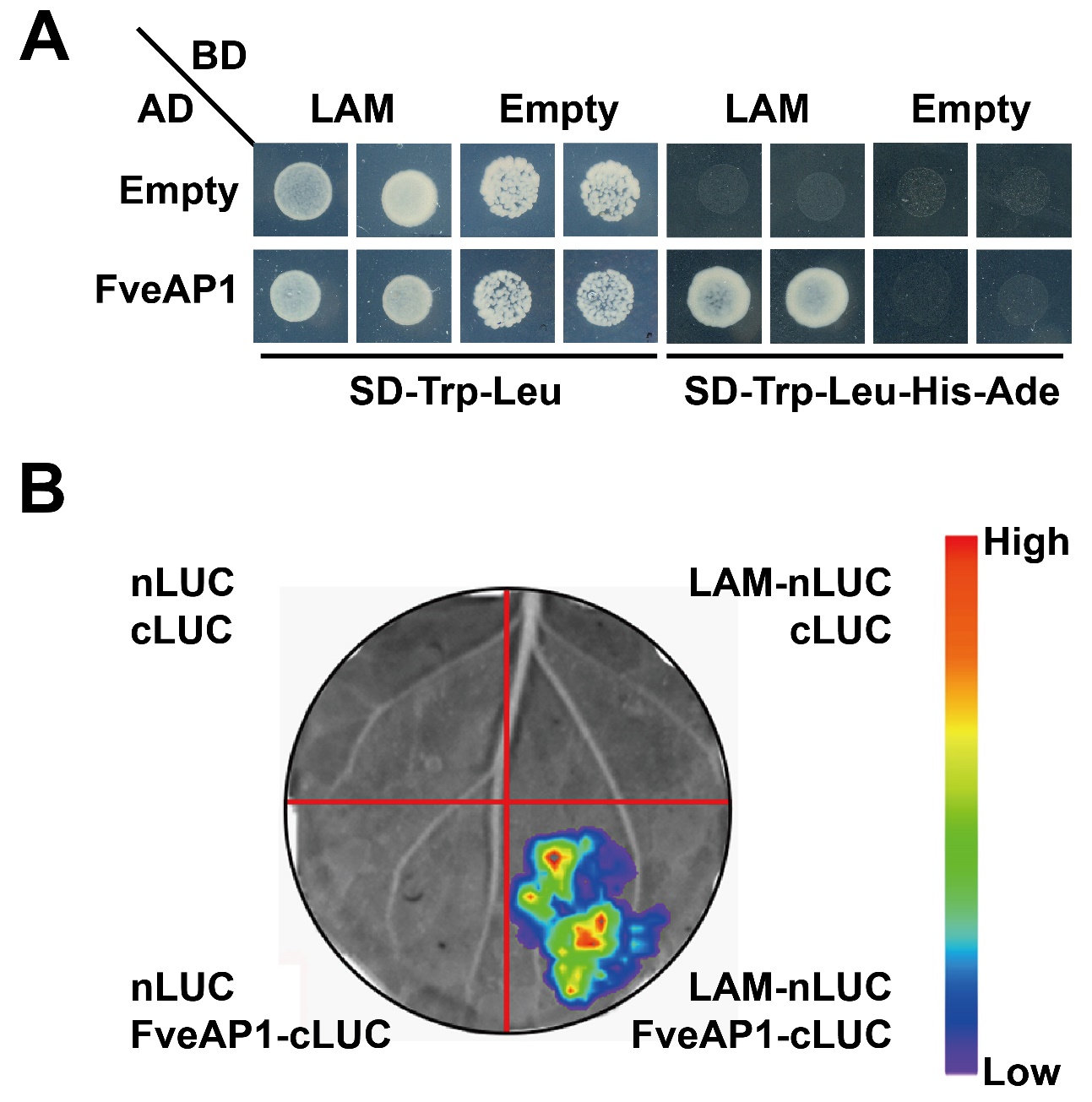

**Supplemental Figure S6. FveAP1 directly interacts with LAM.**

A, FveAP1 interacts with LAM in the Y2H assay. Transformed yeast cells were grown on SD-Leu-Trp and SD-Leu-Trp-His-Ade, respectively. AD, activation domain; BD, DNA-binding domain. Empty vectors were used as controls. B, FveAP1 interacts with LAM in the split luciferase assay. The photograph was taken at 2 days post infiltration into the tobacco leaves.

**Supplemental Table S1. Summary of RNA-seq read statistics.**

| **SampleID** | **Number of raw reads** | **Number of uniquely mapped reads** | **% mapped** |
| --- | --- | --- | --- |
| YW #1 | 53,717,776 | 47,173,870 | 89.84% |
| YW #2 | 50,762,016 | 45,215,072 | 90.99% |
| YW #3 | 56,435,460 | 50,891,634 | 92.14% |
| *fvelfya-1* #1 | 52,948,808 | 47,965,732 | 92.65% |
| *fvelfya-1* #2 | 69,598,176 | 59,493,402 | 87.39% |
| *fvelfya-1* #3 | 49,396,108 | 42,310,494 | 87.54% |
| Average |  |  | 90.09% |

**Supplemental Table S2. Primers used in this study.**

| **Primer Names** | **Primer sequences (5’-3’)** |
| --- | --- |
| **Primers for genotyping** | |
| FveLFYa-F | GCGAGTTTGTTCAAGTGGGA |
| FveLFYa-R  for *fvelfya-1*, *fvelfya-2* | CATCCATAGCGTTGGTGTCA |
| FveLFYa-R  for *fvelfya-3*, *fvelfya-4* | GAAGGGGTGCTCCCTCTGTC |
| FveLFYc-F | CGACTCTCTTCAAGTGGGAC |
| FveLFYc-R | CCGCTTCTTCGCCGCCCTAA |
| LAM-F | CGCTCACGGCCTTATTTCCATTCT |
| LAM-R | GAGTCCAAGGGACTGAGCAAACCTTA |
| **Primers for CRISPR-Cas9** | |
| FveLFYa-DT1-BsF | ATATATGGTCTCGATTGTTCACTGTGAGCACGCTTTGTT |
| FveLFYa-DT1-F0 | TGTTCACTGTGAGCACGCTTTGTTTTAGAGCTAGAAATAGC |
| FveLFYa-DT2-R0 | AACCCGACAGCCCTGTCCAACCCAATCTCTTAGTCGACTCTAC |
| FveLFYa-DT2-BsR | ATTATTGGTCTCGAAACCCGACAGCCCTGTCCAACCCAA |
| FveLFYc-DT1-BsF | ATATATGGTCTCGATTGAGGAGGACTGGCCTCTCTCGTT |
| FveLFYc-DT1-F0 | TGAGGAGGACTGGCCTCTCTCGTTTTAGAGCTAGAAATAGC |
| FveLFYc-DT2-R0 | AACGGGGAGGTGGCACGAGGCACAATCTCTTAGTCGACTCTAC |
| FveLFYc-DT2-BsR | ATTATTGGTCTCGAAACGGGGAGGTGGCACGAGGCACAA |
| **Primers for LFY homologues overexpression** | |
| FveLFYa-ox-F | CTTCACTGTTGATACATATGATGGATCCAAATGCGTTCAC |
| FveLFYa-ox-R | CCCTTGCTCACCATGAATTCGTAGGGAAGCGGATCAGCAG |
| FveLFYb-ox-F | CTTCACTGTTGATACATATGATGGATCCATACTCGATCAC |
| FveLFYb-ox-R | CCCTTGCTCACCATGAATTCGTAGGGCAGCTGATGAGCAG |
| FveLFYc-ox-F | CTTCACTGTTGATACATATGATGGATCCAGACGCGTCCAC |
| FveLFYc-ox-R | CCCTTGCTCACCATGAATTCTGCAGCAGTATCCTTTCCAC |
| FveLFYd-ox-F | CTTCACTGTTGATACATATGATGGTGGGGAGTGATGGAGA |
| FveLFYd-ox-R | CCCTTGCTCACCATGAATTCTGCAGCAGTACCCTTTCCAC |
| **Primers for Y1H assay** | |
| AtLFY-Y1H-F | GATTATGCCTCTCCCGAATTCATGGATCCTGAAGGTTTCACG |
| AtLFY-Y1H-R | AGAAGTCCAAAGCTTCTCGAGCTAGAAACGCAAGTCGTCGC |
| FveLFYa-Y1H-F | GATTATGCCTCTCCCGAATTCATGGATCCAAATGCGTTCAC |
| FveLFYa-Y1H-R | AGAAGTCCAAAGCTTCTCGAGTCAGTAGGGAAGCGGATCAG |
| FveLFYc-Y1H-F | GATTATGCCTCTCCCGAATTCATGGATCCAGACGCGTCCAC |
| FveLFYc-Y1H-R | AGAAGTCCAAAGCTTCTCGAGTCATGCAGCAGTATCCTTTC |
| AtAP1-promoter-F | TTTGATATTGGATCGGAATTCGATTAACAACATAGCACATATTCAACTG |
| AtAP1-promoter-R | ATACAGAGCACATGCCTCGAGACTTCTTACTCTAAAAGAACC |
| FveAP1-promoter-F | TTTGATATTGGATCGGAATTCGCCAGGAAACTGAAATCCGCAC |
| FveAP1-promoter-R | ATACAGAGCACATGCCTCGAGCTACCTGAATCAATGACTACTTTCC |
| AtAG-intron-F | TTTGATATTGGATCGGAATTCCAGTTGTCAACTCTCATGTCCTG |
| AtAG-intron-R | ATACAGAGCACATGCCTCGAGCAAATCGACCACTTGCACAG |
| FveAG-intron-F | TTTGATATTGGATCGGAATTCGTTCTATGTCATGATTCACAG |
| FveAG-intron-R | ATACAGAGCACATGCCTCGAGGGCCATGATCAAAGTTACAG |
| **Primers for Y2H assay** | |
| LAM-Y2H-F | TCAGAGGAGGACCTGCATATGATGGGTTCATTCAATTCTTCT |
| LAM-Y2H-R | TCGACGGATCCCCGGGAATTCGTTCCAAGAAGAAACTGAGAA |
| FveAP1-Y2H-F | GTACCAGATTACGCTCATATGATGGGAAGGGGTAGGGTTCAGCTG |
| FveAP1-Y2H-R | ATGCCCACCCGGGTGGAATTCTGAAGCAAAGCATCCAAGGTGGCA |
| **Primers for split-LUC assay** | |
| LAM-splitLUC-F | ACGGGGGACGAGCTCGGTACCATGGGTTCATTCAATTCTTCT |
| LAM-splitLUC-R | CGCGTACGAGATCTGGTCGACGTTCCAAGAAGAAACTGAGAAA |
| FveAP1-splitLUC-F | TACGCGTCCCGGGGCGGTACCATGGGAAGGGGTAGGGTTCA |
| FveAP1-splitLUC-R | ACGAAAGCTCTGCAGGTCGACTCATGAAGCAAAGCATCCAAG |
| **Primers for qRT-PCR** | |
| FveLFYa-qRT-F | GAGACGACTTGAGGAGGAGG |
| FveLFYa-qRT-R | CTCCTAATCCTCCTCAGCTTC |
| FveLFYb-qRT-F | AACACTGCCAAGGAGCGTGGT |
| FveLFYb-qRT-R | CGCGTAGCAGTGAACGTAATG |
| FveLFYc-qRT-F | GAGACGACGACTTGAGGAGGA |
| FveLFYc-qRT-R | CCGCTTCTTCGCCGCCCTAA |
| FveLFYd-qRT-F | GGTGGGAGAATTCCATCTCTA |
| FveLFYd-qRT-R | AGCAGTGAACGTAATGCCGC |
| AP3a-qRT-F | CGATGAGTACCAGAAGCTTC |
| AP3a-qRT-R | TGATCTGCCTCTTCAGACTC |
| AP3b-qRT-F | CACGGTTCTGTGTGATGCTC |
| AP3b-qRT-R | AGCTCCATAGATCGATCTGC |
| PIa-qRT-F | AACAGGCAGGTGACCTATTC |
| PIa-qRT-R | GCTTGGTATCCCACAACCTC |
| PIb-qRT-F | GAGGTTATGGGATGCAAAGC |
| PIb-qRT-R | ATGTCGTGGAGATTGGGCTG |
| AP1-qRT-F | AACATGCTAGGCTGAAGGTG |
| AP1-qRT-R | CCTCTGTAGCTCAGAGATCG |
| AG-qRT-F | GGATCTGCCTCAGAAGCTAC |
| AG-qRT-R | CCATGCCCTTGAGCTCCTTG |
| FveGAPDH-qRT-F | TCTTTGATGCCAAGGCTGGA |
| FveGAPDH-qRT-R | TCACACGGGAACTGTAACCC |
| ACTIN2-qRT-F | TCCCTCAGCACATTCCAGCA |
| ACTIN2-qRT-R | GATCCCATTCATAAAACCCCAGC |
